# Conserved phenotype and function of human brain border-associated macrophages in iPSC-derived models

**DOI:** 10.64898/2025.12.11.693582

**Authors:** Helena J. Barr, Constanze Depp, Maximilian Hingerl, Trang Nguyen, Justin M. Reimertz, Emma Connolly, Jasmin Patel, Jordan L. Doman, Samuel E. Marsh, Beth Stevens

**Affiliations:** Harvard Division of Medical Sciences, Program in Neuroscience, Boston, MA; Boston Children’s Hospital, Kirby Neurobiology Center, Boston, MA; Broad Institute of MIT and Harvard, Cambridge, MA; Society of Fellows, Harvard University; Howard Hughes Medical Institute

## Abstract

Border-associated macrophages (BAMs) are increasingly implicated in protective brain functions including the removal of pathogenic material such as amyloid-beta (Aβ). However, there is a lack of available and deeply characterized human BAM models. Here, we report that postnatal transplantation of induced pluripotent stem (iPS) cell-derived hematopoietic progenitors to the murine brain sufficiently chimerizes the brain border immune compartment to allow functional interrogations. Human xenotransplanted BAMs (xBAMs) line the leptomeninges and brain vasculature beyond the glia limitans. Via single-cell RNA sequencing, we show that a conserved transcriptional signature distinguishes BAMs from microglia across species origin, age, and genetic background. Both xBAMs and murine BAMs are defined by a hyper-endocytic phenotype and function, surpass other brain macrophages in acute Aβ scavenging, and exhibit compartment-restricted sampling of parenchymal material. Using a modified differentiation protocol, we find that we can generate iPS-derived BAM-like cells (iBAMs) *in vitro*, which are also characterized by a hyper-endocytic phenotype and enhanced engulfment capacity relative to iPS-derived microglia-like cells (iMGLs). Together, these data define a conserved hyper-endocytic BAM phenotype and provide a toolbox for studying human BAMs both *in vivo* and *in vitro*.

## INTRODUCTION

Tissue-resident macrophages populate most mammalian organs and orchestrate key tissue defense and repair functions. In the central nervous system (CNS), tissue-resident macrophages are by far the predominant immune cell type and comprise two anatomically distinct subsets: microglia residing within the brain parenchyma and border-associated macrophages (BAMs) lining the meninges and brain vasculature. Both microglia and BAMs are long-lived cells of embryonic origin distinct from monocyte-derived macrophages (MDMs)^1–4^. BAMs not only exert in classical immune functions such as defense against viral and bacterial pathogens^5,6^, but have also been implicated in clearing pathogenic proteins including amyloid-beta (Aβ)^7,8^. Despite this therapeutic relevance, BAMs are almost exclusively studied in mouse models. Few studies have probed the *in vivo* phenotype and function of human BAMs, in part due to their sparseness in tissue isolates and further because primary human tissues do not allow for experimental manipulation. There is a need for validated models of human BAMs to assess the therapeutic potential of targeting BAMs to treat brain diseases,

Macrophages derived from either circulating monocytes or bone marrow hematopoietic stem cells (HSCs) fail to recapitulate key features of yolk sac-derived tissue-resident macrophage phenotype and function^9^, including in brain engraftment studies^10–12^. The advent of induced pluripotent stem cell (iPSC) technology has instead enabled generation of myriad cell types from somatic cells such as macrophages^13,14^. iPS-derived macrophages replicate key features of microglial biology, both *in vitro* and following *in vivo* engraftment^9,15–20^. Whether iPS-derived macrophages can recapitulate *in vivo* BAM phenotype and function remains underexplored.

Here, we report that early postnatal intracranial transplantation of iPS-derived hematopoietic progenitor cells (HPCs) sufficiently chimerizes the brain border immune compartment to enable functional interrogation of human border-dwelling macrophages in addition to the parenchyma, where cells acquire a microglial fate. Anatomically, xenotransplanted BAMs (xBAMs) line the leptomeninges and brain vasculature of chimeric mice. Single-cell RNA sequencing (scRNAseq) reveals that xBAMs share a core transcriptional signature with murine BAMs across age and murine background strains. We further show that xBAMs exhibit a conserved hyper-endocytic phenotype and are expert scavengers of fluid-borne Aβ, and that this phenotype and function can be replicated *in vitro* using a modified differentiation protocol to generate iPS-derived BAMs (iBAMs). Together, these data provide evidence for a conserved phenotypic and functional profile of human, border-dwelling CNS macrophages and a validated toolkit for assessing their function *in vivo*.

## RESULTS

### Human macrophages establish a brain border immune compartment following xenotransplantation in the mouse

We employed established protocols to generate mouse–human microglial chimeras by transplanting induced pluripotent stem cell (iPSC)–derived hematopoietic progenitor cells (HPCs; WTC11 line) into the brains of immunocompromised *Rag2^-/-^Il2rg^-/-^*mice with additional humanization of *CSF1* to enable engraftment of human hematopoietic cells^16^ (see **Methods, Figure S1A**). Transplantation was performed between postnatal days 2-4. Endogenous microglia were not depleted prior to transplantation as done in other protocols^19,20^. We opted to sufficiently maintain the endogenous myeloid compartment to have a sizable within-individual control for the effect of immunocompromisation on cellular phenotype.

In this approach and similar models, it had been previously noticed that xenotransplanted cells could be found along brain vasculature and within the subdural meninges, and that these cells acquired BAM-like gene expression profiles^16,19^. We hypothesized that iPSC-derived cells within these compartments would take on the phenotype and function of BAMs. To this end, we aged chimeras to adulthood (5 months) and first characterized cell distribution and chimerism via immunohistochemistry. Human macrophages, identified by IBA1 staining in conjunction with a human nuclear marker, tiled the entire murine brain alongside endogenous macrophages (**Figure 1B,C**). Notably, we observed human cells residing within brain border compartments, such as along brain vasculature (**Figure 1C**). We next investigated if these cells may have taken on a BAM-like identity and represent a human xenotransplanted (xBAM) population. To verify this, we stained chimeric brain sections for human CD163, a marker that distinguishes BAMs from microglia across human and murine specimens^4,7,21,22^. CD163^pos^ human macrophages were dispersed across the entire brain, including the leptomeninges (**Figure 1D, Figure S1B**) and along brain vasculature (**Figure 1E**). CD163^pos^ human cells did not colocalize with TMEM119^pos^ cells (**Figure 1E**), indicating successful segregation of BAM-versus microglia-enriched markers.

**Figure 1.**
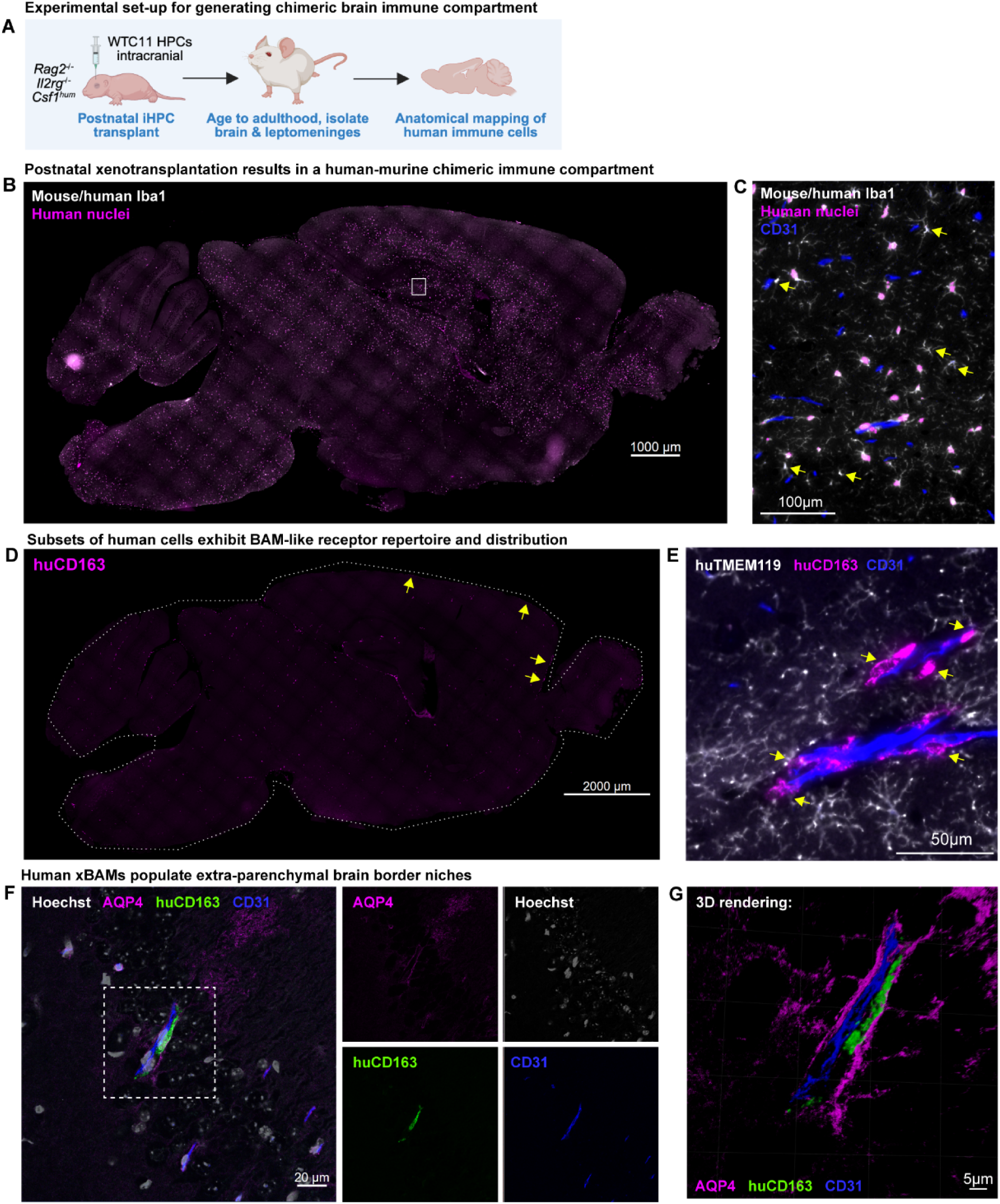
Human macrophages establish a brain border immune compartment in the mouse following xenotransplantation. (**A**) Schematic diagram of experimental approach for generating human-murine chimeric brain immune compartment. (**B and C**) Whole-section tile scan demonstrating distribution of human-derived cells in chimeric brain and zoomed image demonstrating co-habitation of both human (magenta nuclei) and murine macrophages (yellow arrows). (**D**) Whole section tile scan of distribution of human CD163^+^ cells in chimeric brain. Yellow arrows point to example cells localized in leptomeninges. (**E**) Zoomed image demonstrating co-habitation of both human microglia-like cells (xMGs; TMEM119^+^ CD163^-^) and human BAM-like cells (xBAMs; TMEM119^-^ CD163^+^; yellow arrows). (**F and G**) Confocal image and 3-dimensional rendering of xBAM along brain vasculature. Human CD163^+^ xBAM is located beyond glia limitans as evidenced by AQP4 stain. See also **Figure S1**.

To confirm that the putative xBAMs were in fact located in extra-parenchymal brain border niches, we labeled aquaporin-4 (AQP4), a marker of astrocyte endfeet that delineates the glia limitans. Indeed, both perivascular and leptomeningeal xBAMs were situated beyond the glia limitans (**Figure 1F-G, Figure S1B**). Three-dimensional reconstruction of brain vasculature from high-resolution z-stacks demonstrated that the human CD163^pos^ cells were positioned between the abluminal surface of brain vascular endothelial cells and the glia limitans perivascularis (**Figure 1G**), conforming with the anatomical positioning of BAMs within the brain. Taken together, our histological characterization of the chimeric immune compartment revealed that a subset of human cells had taken on the anatomical localization and surface receptor repertoire of *bona fide* BAMs.

### A conserved transcriptional signature distinguishes murine and human BAMs from microglia

To further characterize these human border-resident macrophages, we performed single-cell RNA sequencing of both human and murine immune cells (CD45+) from 3 month-old adult chimeric mice (**Figure 2A**). Because BAMs are relatively sparse compared to microglia, we FACS-enriched both murine and human CD206^hi^ cells prior to sequencing (**Figure S2A**). Samples were treated with transcriptional and translational inhibitors to preserve *in vivo* transcriptional phenotypes^23^. Following doublet removal and quality control, we identified 13,385 immune cells including both human *PTPRC*^+^ (encoding CD45) cells and endogenous murine *Ptprc^+^* cells (**Figure 2B-D, Table S1**). Endogenous mouse cells sampled were primarily microglia and BAMs, as well as a small number of MDMs, classical dendritic cells type 1 and 2 (cDC1/2s), plasmacytoid dendritic cells (pDCs), and neutrophils (**Figure S2B-D, Table S1**). CD45+ human cells were restricted to xenotransplanted microglia and BAM identities (xMGs and xBAMs, respectively), as well as a cluster of proliferating cells (xProlif, **Figure S2E-G, Table S1**).

**Figure 2.**
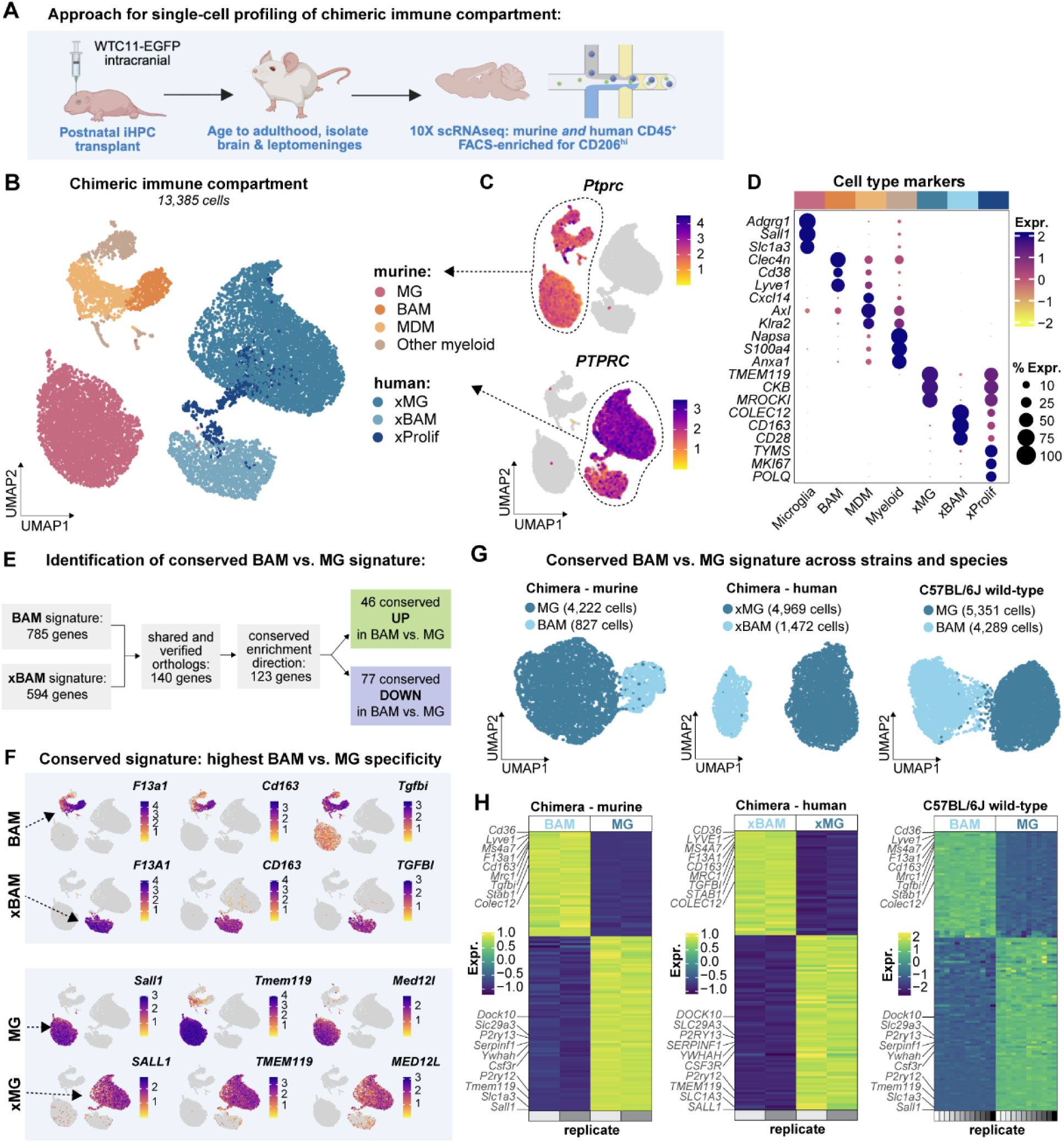
A conserved transcriptional signature distinguishes BAMs from microglia across strains and species origin. (**A**) Schematic diagram of experimental design. Mice received a postnatal transplant of iPS-derived HPCs (WTC11-EGFP). In adulthood, the chimeric brain immune compartment was profiled by scRNAseq enriched for CD206^hi^ cells. (**B-D**) UMAP of annotated cell types, feature plot of human versus murine *Ptprc/PTPRC* expression, and heat map of annotated cell type markers. MG = microglia, BAM = border-associated macrophage, MDM = monocyte-derived macrophage, xMG = xenotransplanted human microglia, xBAM = xenotransplanted human BAM, xProlif = proliferating xenotransplanted human cell (cell type markers in **Table S1**; n of 2 biological replicates). (**E**) Work flow for identification of conserved BAM vs. microglia (MG) signature (full shared signature gene set in **Table S2**). (**F**) Feature plot of expression of top genes distinguishing BAMs/xBAMs from MGs/xMGs. (**G**) UMAPs of datasets used for mapping expression of conserved signature gene set: murine MGs and BAMs from the chimeras (n of 2 biological replicates), human MGs and BAMs from the chimeras (n of 2 biological replicates), and MGs and BAMs from C57BL/6J mice (aged 2.5 months; all males; n of 12 biological replicates). (**H**) Heatmap of relative expression of conserved signature gene set across datasets and cell types. See also **Figure S2**.

We then sought to determine conserved markers distinguishing BAMs from microglia across species. We started with cell-level differential gene expression (DEG) analysis, separately per species, followed by orthology verification on DEGs (**Figure 2E, Methods**). We identified a set of 123 genes that formed a conserved BAM versus microglia signature across human and murine cells (**Table S2**). Of this conserved gene set, we first examined those with the highest specificity for each cell type (**Figure S2H**). BAMs and xBAMs were highly enriched for the coagulation factor XIII (*F13a1/F13A1)*, the scavenger receptor CD163 (*Cd163/CD163)*, and the transforming growth factor beta (TGFβ)-induced gene *Tgfbi/TGFBI* (**Figure 2F**). *TGFBI* encodes a protein that binds to both type I and type IV collagens^24^, which are major leptomeningeal and perivascular structural components and consistent with the brain border localization of xBAMs^25–27^. Microglia and xMGs were highly enriched for the transcription factor *Sall1/SALL1*, the transmembrane protein *Tmem119/TMEM119*, and the mediator complex subunit *Med12l/MED12L* (**Figure 2F**). These highest-specificity genes from the conserved gene set have all been reported as BAM- or microglia-enriched in sequencing studies of both human and murine brains^21,22,28–30^.

We then validated the conserved microglia versus BAM gene set by probing its expression across multiple datasets. As expected, the signature obtained from the chimeras was highly distinctive of chimeric BAMs and microglia of both murine and human origin (**Figure 2G-H**). However, it also effectively distinguished BAMs and microglia from healthy, wild-type, young adult, brains from C57BL/6 mice (**Figure 2G-H**), as well as BAMs and microglia from the brains of aged wild-type C57BL/6 mice (**Figure S2I**). Given evidence that the transcriptional phenotype of brain immune cells varies around the day-night cycle^31^, we also examined whether the conserved gene set could distinguish between microglia and BAMs around the clock. K-means clustering of samples obtained at Zeitgeber Times (ZT, corresponding to the number of hours elapsed since the beginning of the light cycle) ZT0, ZT6, ZT12, and ZT18 successfully distinguished microglia and BAMs at all four time points using only the conserved gene set (**Figure S2J**). Taken together, these data indicate that the conserved transcriptional signature distinguishing human and murine BAMs from microglia was also discriminatory across time, age, and murine background strain.

### Human and murine BAMs are defined by a hyper-endocytic phenotype and function

To explore the defining features of BAMs versus microglia, we performed Ingenuity Pathway Analysis (IPA) on the conserved gene set to identify cellular processes enriched in BAMs. Many of the top pathways were involved in engulfment including clathrin-mediated endocytosis (CME) and the binding and uptake of ligands by scavenger receptors (**Figure 3A**). Indeed, across both the murine and human immune compartments of chimeric mice, as well as in re-analysis of thousands of BAMs and microglia from C57BL/6J mice, CME-associated genes were massively enriched in BAMs and xBAMs (**Figure 3B**). This included genes encoding receptors known to be internalized via CME (*CD163, CD206, STAB1*) as well as components of the clathrin vesicles (*CLTC*), proteins involved in initial CME nucleation (I*TSN1, EPS15*), and cargo selection and accessory proteins (*DAB2, AP2A2, AP2S1*). Thus, human and murine BAMs are characterized by a hyper-endocytic transcriptional phenotype.

**Figure 3.**
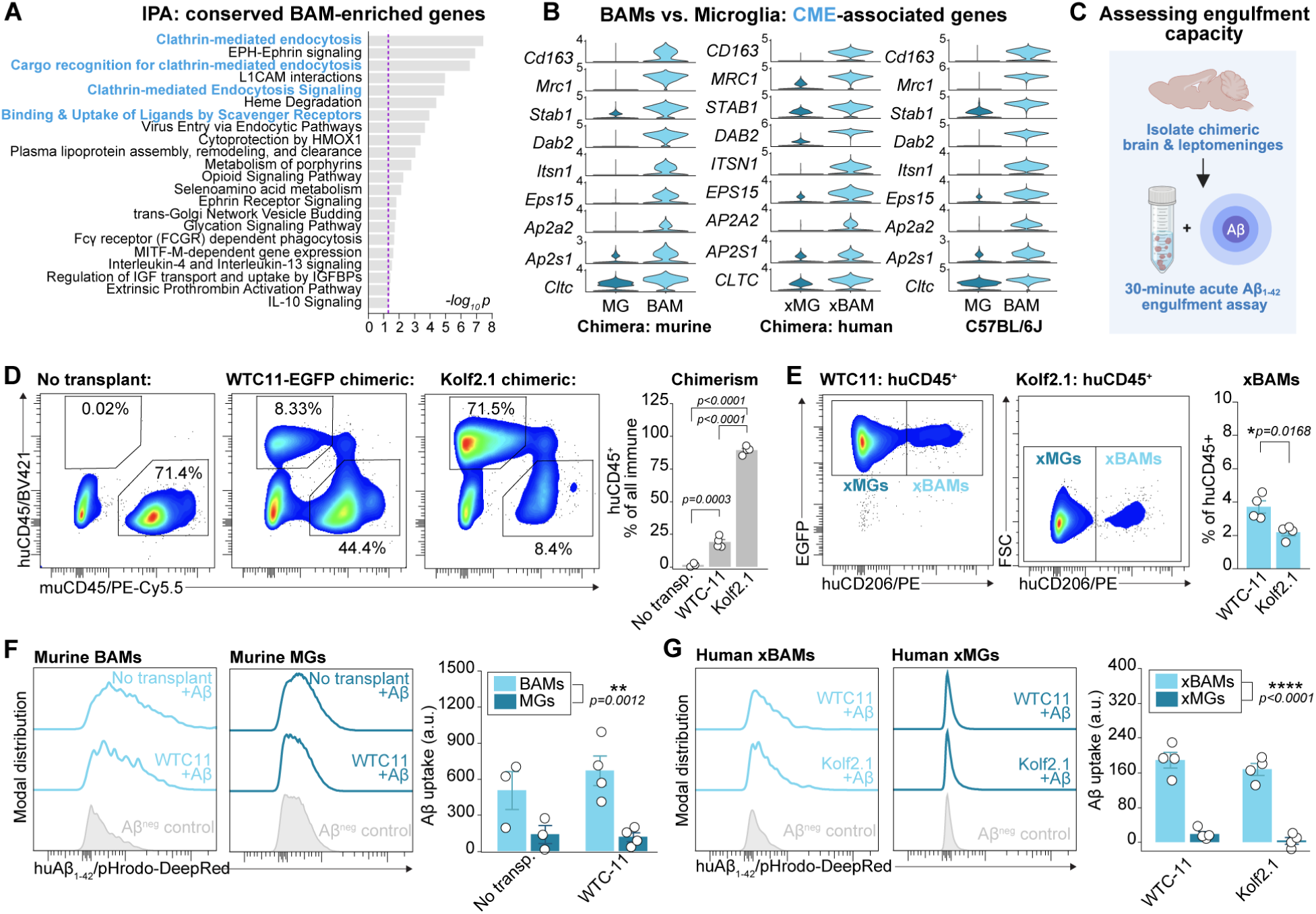
Human and murine BAMs are defined by a hyper-endocytic phenotype and function. (**A**) IPA on genes enriched in BAMs relative to microglia in conserved gene set. Dashed line represents significance threshold. (**B**) Violin plot of expression of core CME-associated genes in BAMs versus microglia of murine and human origin in chimeras, as well as BAMs versus microglia from wild-type C57BL/6J mice. (**C**) Experimental design of acute *ex vivo* engulfment assay for testing scavenging capacity. (**D**) Representative pseudo-colored plots of murine CD45 (muCD45) and human CD45 (huCD45) expression in untransplanted, WTC11-EGFP-transplanted, and Kolf2.1-transplanted chimeras. Double-positive cells are likely doublets. Chimerism (% of all CD45^+^ cells that are huCD45^+^) varies by condition (F_2,8_=737.96, p<0.0001; one-way ANOVA with Tukey’s HSD post-hoc testing; n of 3-4 independent replicates per condition). (**E**) Representative pseudo-colored plots of human CD206 (huCD206) expression in huCD45^+^ cells from WTC11-EGFP-transplanted and Kolf2.1-transplanted chimeras. A slightly higher proportion of huCD45^+^ were xBAMs in WTC11-EGFP versus Kolf2.1-transplanted chimeras (two-tailed student’s t-test; n of 4 independent replicates per condition). (**F**) Representative histograms and group-level quantification of pHrodo-DeepRed conjugated Aβ_1-42_ fibril uptake by murine BAMs and microglia across untransplanted and WTC11-EGFP-transplanted mice. Main effect of cell type (p=0.0012) but not transplant group (p=0.6106), and no interaction effect between the two (p=0.2379) on Aβ uptake (mixed effects model accounting for variance due to individual sample; n of 3-4 independent replicates per condition). Note: Kolf2.1-transplanted mice were not included in this comparison due to human-dominance of chimeric compartment (<100 murine BAMs detected in some samples). (**G**) Representative histograms and group-level quantification of pHrodo-DeepRed conjugated Aβ_1-42_ fibril uptake by human xBAMs and xMGs across WTC11-EGFP-transplanted and Kolf2.1-transplanted mice. Main effect of cell type (p<0.0001) but not transplant group (p=0.2743), and no interaction effect between the two (p=0.7315) on Aβ uptake (mixed effects model accounting for variance due to individual sample; n of 4 independent replicates per condition). *For all panels: points represent individual mice, bars and error bars represent mean and SEM. Normalization to background is done per cell type with cell type-matched negative control to account for cell type differences in background autofluorescence*. See also **Figure S3**.

To interrogate whether the hyper-endocytic phenotype of BAMs and xBAMs was functionally relevant, we designed acute engulfment assays where chimeric brains were acutely dissected and digested for 30 minutes within a fixed concentration of exogenous, pHrodo-conjugated Aβ_1-42_ preformed fibrils (**Figure 3C**). This timing precluded prolonged phagocytic uptake and ensured all samples saw an equal concentration of substrate for an equal duration. Aβ uptake was subsequently measured on the single-cell level via flow cytometry using a validated panel for identifying xBAMs (**Figure S3, Table S3**). We performed these assays using two different models: WTC11-EGFP- and Kolf2.1J-transplanted chimerics, representing two very commonly used iPSC lines. In both cases, a chimeric immune compartment was established including a robust xBAM population (**Figure 3D-E**), though the rate of chimerism was higher in Kolf2.1J chimeras. Across both transplanted and un-transplanted mice, murine BAMs engulfed far more Aβ relative to microglia (**Figure 3F**). Aβ signal was on average 5-fold higher in BAMs than microglia. This effect was also observed in human cells: xBAMs from both WTC11-EGFP- and Kolf2.1J-transplanted chimerics by far exceeded xMGs in Aβ uptake (**Figure 3G**). Aβ signal was on average 16-fold higher in xBAMs than xMGs. xMGs performed poorly in the acute assays, exhibiting little uptake detectable beyond background autofluorescence. Together, these data indicate that human and murine BAMs are distinguished by a hyper-endocytic functional phenotype.

### Validating toolkits for assessing human macrophage function *in vivo*

We next validated a toolkit to measure *in vivo* engulfment by brain macrophages, FEAST, in the chimeric mice. FEAST is a flow cytometry pipeline that abrogates *ex vivo* artifacts through the combination of 1) perfusion fixation prior to tissue dissociation, and 2) a cellular acid wash to detach debris bound to the cell surface, prior to 3) permeabilization and labeling of engulfed substrates^32^. As a positive control, we fed adult chimeras a diet of 0.2% Cuprizone (Teklad) for a duration of four weeks to induce demyelination (**Figure S3A**). Using FEAST, we had previously found that microglia robustly respond to demyelination by engulfing myelin basic protein (MBP), but that no such response is seen in BAMs^32^. Following the fixed-cell FEAST protocol, both human and murine immune cells were detectable via flow cytometry and cuprizone administration did not alter chimerism nor the percent of human cells that were xBAMs (**Figure S3B**). Cuprizone-mediated demyelination led to a large increase in MBP engulfment by both microglia and xMGs (**Figure S3D-F**). Even in the absence of cuprizone, xMGs were enriched for MBP signal relative to endogenous microglia (**Figure S3F**). As previously observed, little to no MBP was detectable above background in either BAMs or xBAMs, and cuprizone demyelination had no impact on this signal (**Figure S3G-H**). Thus, as in C57BL/6 mice, xBAMs maintained compartment-specific sampling of brain-enriched proteins.

For both *in vivo* engulfment assays using FEAST and acute *ex vivo* engulfment assays, we identified recommended gating strategies and validated human-specific antibodies (**Figure S3I-L**). We found that, in chimeric mice, both the anti-human-CD45 and anti-human-CD206 show high specificity against negative and isotype controls (**Figure S3B, Figure S3L**). We also found that all human cells in chimeric mice, detected via flow cytometry, had taken on a CD45^+^ phenotype, and that all CD45^+^ cells were additionally CD11b^+^ (**Figure 3J-K**). Together, these data demonstrate that xenotransplanted cells in human-mouse chimerics can be phenotyped and functionally probed using validated antibodies and gating strategies.

### A modified differentiation protocol generates iPS-derived BAM-like cells *in vitro*

We next sought to determine whether we could generate human iPS-derived BAM-like cells *in vitro* for future high-throughput applications. A large body of evidence indicates that TGF-β drives microglial identity^2,4,12,33–39^, and protocols for *in vitro* generation of human iPS-derived microglia-like cells (iMGLs) that most closely resemble human microglia supplement the differentiation media with TGFβ1^17^. Microglia lacking SMAD4, a central protein downstream of TGF-β signaling, also exhibit an immature phenotype and upregulated endocytic receptors such as CD206, whereas BAM phenotype is unaffected by SMAD4 deletion^4^. Thus, we posited that altering TGF-β exposure may shift the identity of iPS-derived macrophages to a BAM-like phenotype rather than microglial. Thus, we simultaneously differentiated the same batch of WTC11-EGFP iHPCs to iMGLs either with or without TGFβ1 supplementation (“iMGL” versus “iBAM” protocol) (**Figure 4A**), and then performed bulk-RNA sequencing and engulfment assays. Obvious morphological differences were observed upon TGFβ1 omission when cells were cultured on Matrigel-coated plates (**Figure 4B**). Cells from the iBAM condition acquired a larger, elongated shape and were filled with vesicular structures, whereas classical iMGLs were smaller, ramified, and possessed fine cellular processes, reminiscent of the morphological differences between BAMs and microglia *in vivo* (**Figure 1E**). Transcriptionally, genes enriched in the iBAM condition were frequently top BAM genes from the conserved BAM versus microglia gene set (*F13A1, DAB2, MRC1, COLEC12, LYVE1*), whereas genes enriched in the iMGL condition were from the core microglial signature (*CX3CR1, YWHAH, SERPINF1, P2RY12*; **Figure 4C-E**). This was further supported by the strong linear correlation between *in vitro* and *in vivo* transcriptional phenotypes whereby cells from the iBAM condition closely resembled xBAMs and those from the iMGL condition closely resembled xMGs (**Figure 4F**). Together, these data indicate that omission of TGFβ1 yields iPSC-derived macrophages that more closely resemble the transcriptional identity of BAMs than microglia.

**Figure 4.**
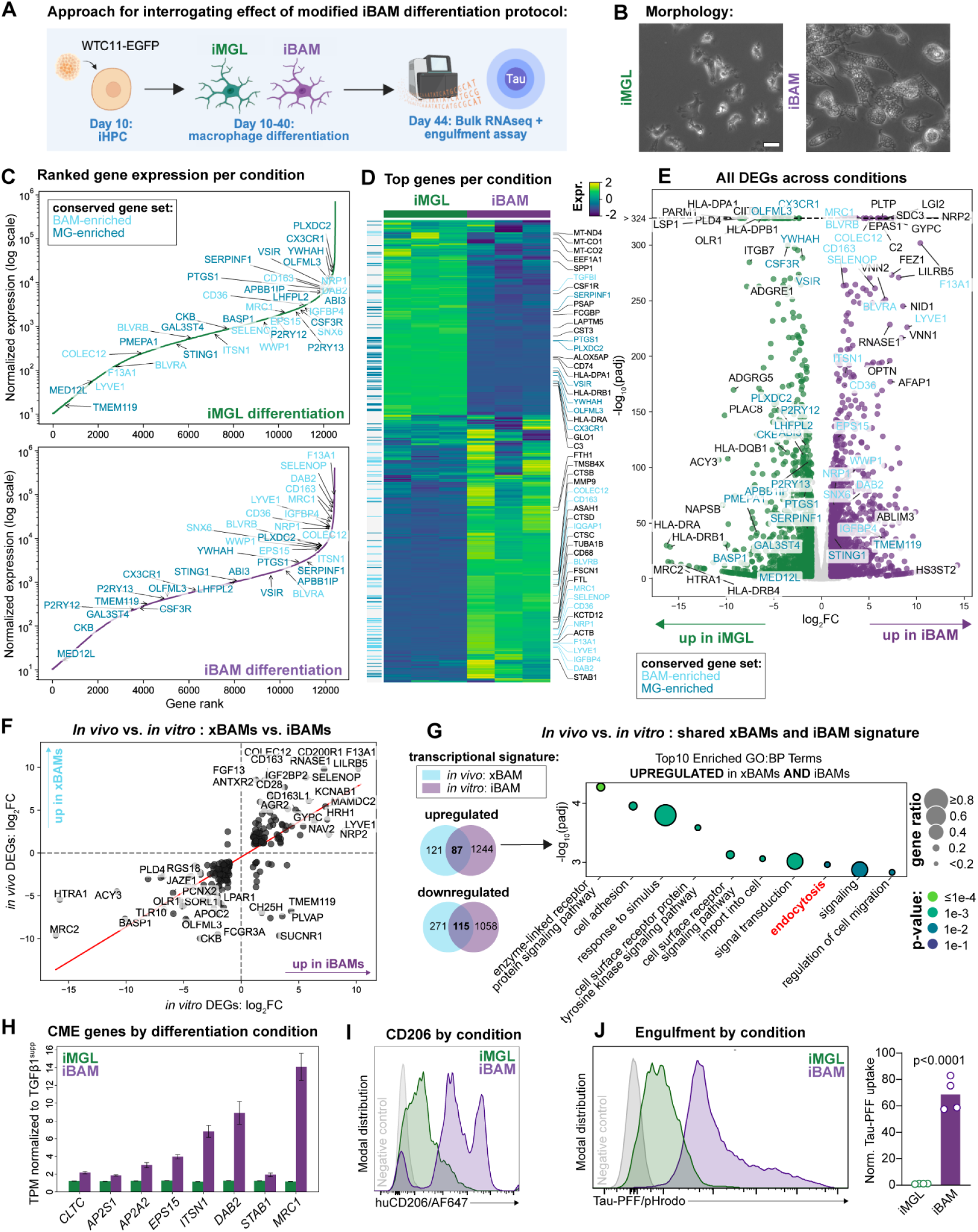
A modified differentiation protocol generates iPS-derived BAM-like cells in vitro. (**A**) Experimental setup to study the effect of a modified differentiation protocol on iPS-derived macrophage phenotype *in vitro*. iHPCs were subjected to either a standard iMGL differentiation or a modified “iBAM” differentiation protocol starting at Day 10. Transcriptional profiles were assessed using bulk-RNA sequencing and engulfment capacities were assessed using flow cytometry. (**B**) Representative images of cell morphology towards the end of the differentiation (day 36). Note the size difference as well as vesicular structures emerging in iBAMs vs. iMGLs. (**C**) Ranked gene expression plots for iMGLs vs iBAMs (n of 3 per condition, with n representing separate wells from the same differentiation). Only genes with a TPM > 10 are plotted. Labeled genes are from the conserved BAM vs. microglia gene set, as well as their enrichment direction.(**D**) Heatmap showing the expression of the 916 most abundant genes (top 2.5% per culture type) across samples. Expression values for each gene are Z-score scaled across samples. The vertical color bar on the left indicates genes from the conserved BAM vs. microglia gene set, as well as their enrichment direction. (**E**) Volcano plot comparing gene expression in iMGLs vs iBAMs. Significantly regulated genes (padj < 0.01, |log2FC| > 1) are colored. The 20 most significant genes on both the up- and down-regulated sides are labeled, along with key marker genes. (**F**) Scatter plot correlating the log2 fold changes of DEGs from an xBAM versus xMG DEG analysis, against DEGs from the iMGL vs. iBAM DEG analysis. The 40 most regulated genes in this common set are labeled. (**G**) Venn diagrams illustrating the overlap of DEGs from the xBAM versus xMG DEG analysis, and DEGs from the iMGL vs. iBAM DEG analysis, and Gene Ontology (GO) enrichment analysis for the overlapping set of genes identified as commonly regulated between xBAMs and iBAMs. (**H and I**) Bar plot displaying expression of eight endocytosis-related genes of interest per condition and histogram showing protein-level upregulation of CD206 in the iBAM condition via flow cyometry. For bar plot, error bars indicate the standard error of the mean (SEM) between replicates, n of 3 replicates per condition. (**K**) Representative histogram and quantification of pHrodo-labelled Tau-PFF uptake by iMGLs versus iBAMs. Bar graphs show mean fluorescence intensity of pHrodo-signal normalized to control condition. Dots represent single wells (n=4 per condition, 1 differentiation). For statistical analysis, unpaired Student’s t-test was performed.

We next examined whether cells generated using the iBAM protocol could mirror the hyper-endocytic phenotype of BAMs. To do this, we obtained a shared gene set defining both xBAMs and iBAMs; endocytosis was a significantly enriched GO term in this gene set (**Figure 4G**). In fact, many of the top CME-associated genes defining both murine and human BAMs were among the DEGs enriched in the iBAMs (*CLTC, AP2S1, AP2A2, EPS15, ITSN1, DAB2, STAB1*; **Figure H**). Notably, we observed approximately a 14-fold upregulation of *MRC1* in the iBAMs that was reflected on the protein-level (**Figure 4H,I**). To test the functional relevance of this hyper-endocytic signature in iBAMs, we performed engulfment assays using pHrodo-conjugated tau preformed fibrils (PFFs) – a substrate relevant to AD and known to be uptaken by CME^40^. iBAMs exhibited massively increased tau uptake (**Figure 4J**) and engulfed approximately 60 times more in the same amount of time relative to the iMGLs. Thus, we find that an adapted differentiation protocol can generate hyper-endocytic iPS-derived macrophages sharing both transcriptional and functional features with BAMs *in vitro*.

## DISCUSSION

BAMs are a fascinating tissue-resident macrophage population that, despite relative sparseness in the CNS, have been implicated in crucial functions including protection against foreign pathogens, maintenance of vascular homeostasis, and support of fluid flow through the brain and its borders^5–7,41,42^. Despite growing research interest, there is a lack of validated human models for studying BAMs. Here, we demonstrate that chimeric human-mouse models can be used to generate human BAMs possessing a conserved *in vivo* transcriptional phenotype. Furthermore, we validate a series of tools to probe human BAM-like cell functionality *in vivo* and *in vitro*.

Using these models, we show that both murine and human BAMs are defined by a hyper-endocytic phenotype and an exceptional capacity for engulfment of extracellular, fluid-born substrates. Historically, BAMs had been identified in the 1980s and 90s due to their remarkable scavenging of substrates delivered into the brain^43,44^. We propose that the heightened scavenging capacity of BAMs may be driven by their specialized capacity for CME, an engulfment modality that can internalize extracellular substrates on the order of minutes. What do BAMs engulf *in vivo,* and why might BAM engulfment matter in brain homeostasis and disease? Broad BAM depletion studies using clodronate-laden liposomes (CLLs) have implicated BAMs in the clearance of pathogenic brain peptides such as Aβ^7,8^, as their loss precipitates amyloid pathology. Future studies interrogating the functional relevance of BAM engulfment, as well as experimentally boosting this function, may nominate therapeutically-relevant targets for the treatment of disorders characterized by Aβ accumulation, such as Alzheimer’s Disease (AD) and cerebral amyloid angiopathy (CAA).

TGF-β and downstream signaling proteins have been repeatedly implicated in driving microglial identity^2,4,12,33–39^. For this reason, a number of established iMGL protocols supplement the differentiation media with TGFβ1 to obtain cells most closely resembling human microglia^17,45^. We find that TGFβ1 drives downregulation of endocytic phenotype and function in iMGLs, consistent with endocytic phenotype being a major cellular distinction between BAMs and microglia. The cellular mechanisms involved in driving the hyper-endocytic phenotype of BAMs remain less clear. One possibility is that this is a default macrophage signature that is specifically downregulated with the adoption of the microglial fate. A second possibility is that currently uncharacterized mediators drive the hyper-endocytic phenotype of BAMs. Further research, for example using CRISPR screens in iPS-derived HPCs, may uncover novel mechanisms involved in the adoption of BAM versus microglia identity.

In addition, the clear effect of TGFβ1 supplementation on both iMGL phenotype and function indicates that iMGL differentiation protocols lacking TGFβ1 supplementation may be generating cells more akin to BAMs than microglia. Our morphological and transcriptional observations are in line with earlier reports examining the effect of TGFB1 omission from iMGL differentiation protocols leading to robust upregulation of more classical macrophage markers and downregulation of microglia identity markers^45^. This is an important consideration for the interpretation of *in vitro* studies of iPSC-derived microglia-like cells. In the future, it would also be interesting to directly compare the described iBAM protocol to other protocols used to generate iPSC-derived tissue-resident macrophages from other organs^46^.

Another noteworthy observation of the chimeric mouse model is that it further confirms the long-lived nature of BAMs, which has been supported by a large body of evidence indicating that BAMs are yolk sac-derived macrophages sharing a common embryonic origin to microglia^2–4,47^. Indeed, we find that human iPS-derived HPCs delivered to the brain in early development adopt a BAM-like identity and persist for many months (> 5 months) in brain border niches. The existence of xBAMs in chimeric models had been previously reported^16,20^, suggesting that this is a replicable phenomenon across laboratories and experimenters. To our knowledge, we are the first to provide in-depth transcriptional and functional characterization of these human xBAMs. Furthermore, we provide a validated flow cytometry toolset to enrich BAMs and probe their functionality, namely their engulfment capacity. Together, these tools could be applied to interrogate the drivers of BAM identity in the developing brain in addition to therapeutically-relevant applications. For example, an interesting future application would be developing chimeric models possessing only xBAMs and no xMGs to specifically interrogate brain border immune function.

Our study has some limitations. The generation of human-murine chimeras *de facto* necessitates an immunocompromised background strain. Thus, BAMs and microglia developed and persisted in an organism effectively lacking an adaptive immune system. This is impossible to circumvent in the context of xenotransplantation models. However, we performed our analysis such that 1) we could identify and control for effects of immunocompromised status by directly comparing human to murine signatures within the chimeric mice, and 2) we could verify the validity of our conserved BAM signature by probing existing scRNAseq datasets of thousands of BAMs and microglia from C57BL/6 mice. This approach established that our conserved gene set was discriminatory for BAMs and microglia in young and aged immunocompetent, wild-type mice. A second consideration in this study is the iPS-derived nature of the human brain macrophages. As evidenced by the scRNAseq, a small but significant proportion of the human macrophages were in an actively proliferating state that is rare in endogenous murine macrophages from the adult brain. It is unclear if this stems either 1) from the iPSC origin of transplanted human cells or 2) is a consequence of species differences in the maturation timescale of microglia/macrophages. Our scRNA-seq experiment was performed at 3 months of age; transplanted cells are essentially “very young” cells on the scale of human maturation and aging. This caveat might also extend to the non-proliferating xMGs and xBAMs. This question of iPS artifact versus species lifespan difference could be addressed in the future by interrogating transplanted murine iPSC-derived HPCs. Furthermore, it will be interesting to profile xMGs and xBAMs from older and aged animals to understand how the developmental trajectory of human xMGs/BAMs shifts over time.

Together, our work provides systematic characterization of human iPS-derived xenotransplanted BAMs in a chimeric mouse model. We find that, across species and lifespan, BAMs are defined by a hyper-endocytic signature and exceed parenchyma-resident microglia in acute scavenging capacity. We hope that our validation efforts will support future research interrogating the development, function, and therapeutic relevance of BAMs.

## MATERIALS AND METHODS

### Mice

For chimeric generation, an immunocompromised CSF1 humanized strain was used which enables chimerism and the maintenance of human macrophages. Founder animals were purchased from the Jackson Laboratory (C;129S4-Rag2tm1.1Flv Csf1tm1(CSF1)Flv Il2rgtm1.1Flv/J; Jax #017708)^48^, and a colony was established in the animal core at the Broad Institute of MIT and Harvard. All experiments were performed following protocols approved by the Institutional Animal Care and Use Committee (IACUC) of the Broad Institute of MIT and Harvard and in compliance with NIH guidelines for the humane use and treatment of laboratory animals. Mice were group-housed in Sealsafe Plus cages (Techniplast #GM500) with *ad libitum* access to food and water in standard 12-hour-light, 12-hour-dark cycles following all temperature, humidity, and capacity recommendations from the American Association for Accreditation of Laboratory Animal Care (AALAC). Males and females were both used in this study. A subset of mice underwent cuprizone-mediated demyelination, in which case their regular chow was replaced with a 0.2% Cuprizone diet (Teklad #TD.140803) for a duration of four weeks.

### *In vitro* differentiations

#### Stem cell culture

All work involving stem cells underwent review and approval from the Broad Institute Office of Research Subject Protection (OSRP; NHSR-5206). Work with chimeric samples also underwent review and approval from the Boston Children’s Hospital Institutional Review Board (IRB; IRB-P00050348). Induced pluripotent stem cells (iPSCs) were cultured in Essential 8 medium (Thermo Fisher Scientific, #A1517001) on 6-well plates coated with Matrigel (Corning, #356231) with daily media change. Cells were passaged as aggregates every 4-5 days using ReleSR (STEMCELL Technologies, cat #100-0483). In this manuscript we used three iPSC lines: Non-modified WTC11, the WTC11 derivative WTC11-EGFP and Kolf2.1J. The EGFP-WTC11 iPSC line was obtained from Matthew Blurton-Jones^49^. The Kolf2.1J line was obtained from JAX. The WTC-11 line was obtained from the Coriell Institute for Medical Research.

#### iMGL generation

We followed published protocols to differentiate iMGL from iPSC lines first differentiating them to HPCs with slight modifications^15,17,18^. For example, we utilized the embryoid bodies (EBs) method to generate hematopoietic progenitor cells (iHPCs). On day-1, when the iPSCs colonies reached 80%-90% confluency, they were dissociated into a single-cell suspension using Accutase (STEMCELL Technologies, #07920), centrifuged at 300g for 5 minutes and resuspended in Essential 8 with 10μM Y-27632 ROCK inhibitor (Selleckchem, #s1049). Cells were seeded on ultralow adherence six-well plates (Corning, #29443-030) at a density of 200,000 cells per well with 2 mL media for 24 hours in a normoxic incubator (20% O2, 37°C) to form EBs.

For the next 10 days, the EBs were grown in filtered iHPC base medium composed of 50% IMDM (Thermo Fisher Scientific, cat #12440053), 50% F12 (Thermo Fisher Scientific, cat #11765054), 2% v/v Insulin-Transferrin-Selenium-Ethanolamine (Thermo Fisher Scientific, cat #51500-056), 400 μM monothioglycerol (Sigma, cat #M1753), 64 μg/mL L-ascorbic acid 2-phosphate (Sigma, cat #A8960), 10μg/mL polyvinyl alcohol (Sigma, cat #P8136), 1X GlutaMAX (Thermo Fisher Scientific, cat #35050061), 1X chemically defined lipid concentrate (Thermo Fisher Scientific, cat #11905031) and 1X nonessential amino acids (Thermo Fisher Scientific, cat #11140050). On day 0, cells were gently collected using a serological pipette, spun down at 100g for 5 minutes and resuspended in iHPC media supplemented with 12.5ng/mL Activin-A (Thermo Fisher Scientific, cat #PHC9564), 50ng/mL BMP4 (Thermo Fisher Scientific, cat #PHC9534), 50ng/mL FGF2 (Thermo Fisher Scientific, cat #PHG0261), 2mM LiCL (Sigma, cat #L7026), and 1 μM ROCK inhibitor. Cells were cultured under hypoxic conditions (5%O2, 5%CO2, 37°C). On day 2, cells were replated as described on day 0 with iHPC media containing 50ng/mL FGF2 and 50ng/mL VEGF (Peprotech, cat #100-20) and then placed in the hypoxic incubator. On day 4, cells were replated again with iHPC media supplemented with 50ng/mL FGF2, 50ng/mL VEGF, 10ng/mL SCF (Thermo Fisher Scientific, cat #PHC2116),10ng/mL IL-3 (Peprotech, cat #200-03), 50ng/mL IL-6 (Peprotech, cat #AF-200-06), and 50ng/mL TPO (Peprotech, cat #300-18) before being transferred back to a normoxic incubator. On day 6 and 8, 1mL of day 4 medium was added per well. On day 10, the supernatant containing floating iHPCs were filtered from the EBs using a 40μm cell filter, centrifuged at 300g for 5 minutes and either cryopreserved with CryoStor CS10 (STEMCELL Technologies, cat #07930) or continued with iMGL differentiation.

Day 10 iHPCs can be thawed from frozen stock or freshly seeded on Matrigel-coated 6-well plates at a density of 100,000 cells/well with 2mL of iMGL differentiation media containing of DMEM/F12 (Thermo Fisher Scientific, cat #11330032), 2% v/v Insulin-Transferrin-Selenium (Thermo Fisher Scientific, cat #41400045), 2% v/v B27 (Thermo Fisher Scientific, cat #17504044), 0.5% v/v N2 (Thermo Fisher Scientific, cat #17502048), 5μg/mL insulin (Sigma, cat #I2643), 200 μM monothioglycerol (Sigma, cat #), 1X GlutaMAX, and 1X nonessential amino acids supplemented with 25ng/mL M-CSF (Peprotech, cat #AF-300-25), 50ng/mL TGFβ-1 (Peprotech, cat #100-21) and 100ng/mL IL-34 (BioLegend, cat #577906) to the end of the differentiation. Cells were fed with 1mL media every other day. On day 22, floating cells were collected and the wells were incubated with cold DPBS (Sigma, cat #D8537) for 5 minutes and then scraped to detach the adherent cells. All cells were spun down at 300g for 5 minutes and replated at 150,000 cells/well using half of old collected media and half of new iMGL differentiation media. On day 30, cells were replated as day 22 with additional 100ng/mL CD200 (Bon Opus Biosciences, cat #BP004) and 100ng/mL CX3CL1 (Peprotech, cat #300-31). iMGLs were considered mature on day 40 and ready to be used for downstream experiments.

#### Flow cytometric quality control of differentiation

The expression of hematopoietic progenitor cell markers as well as mature microglia markers was assessed on day 10 and day 40 of differentiation in culture, respectively. All cell preparation steps were conducted on ice to maintain cell viability. For each staining condition, 30,000–50,000 cells were seeded into individual wells of a 96-well U-bottom plate. The total volume in each well was adjusted to 200 µL with Dulbecco’s Phosphate-Buffered Saline (DPBS). The plate was centrifuged at 1000×g for 5 minutes to pellet the cells. The supernatant was discarded, and cell pellets were resuspended in 100 µL of a blocking and viability staining solution, which consisted of DPBS with 0.5 mM EDTA, a 1:100 dilution of human Fc-Block reagent (#564220, BD Biosciences) and a 1:1000 dilution of a viability dye (LIVE/DEAD™ Fixable Near IR (876) Viability Kit, L34980, Thermofisher Scientific). The plate was incubated for 10 minutes at 4C, protected from light. Following incubation, cells were centrifuged at 1000×g for 5 minutes, and the supernatant was discarded. The cells were washed once by resuspending the pellet in 200 µL of FACS buffer consisting of DPBS with 0.5 mM EDTA and 1% BSA (Miltenyi Biotech 30-091-376). After another centrifugation step (1000×g, 5 minutes), the supernatant was discarded and cells were resuspended in 100 µL of FACS buffer containing the appropriate antibody panel. For day 10 differentiation of HPCs, single stainings were performed using: anti human CD45 BV421 (304032, Biolegend), APC anti-human CD43 Antibody (343206, Biolegend), PE anti-human CD41 Antibody (303705, Biolegend). Differentiation was determined successful if 30 – 50% of cells showed CD45 positivity and all cells were positive for CD43 and CD41. For day 40 differentiation QC for iMGL stainings were performed using: anti human CD45 BV421 (304032, Biolegend), anti human CX3CR1 Alexa647, anti human CD11B-Alexa 488 (301318) Biolegend, anti human P2RY12 PE (392104, Biolegend). Differentiation was considered successful if all cells were positive for all markers. The cells were incubated with the antibodies for 30 minutes at 4C on a shaker in the dark. After antibody incubation, the cells were washed twice with 200 µL of FACS buffer, with a centrifugation step (1000×g, 5 minutes) following each wash. Samples were analyzed using a CytoFLEX LX cytometer.

### Generation of chimeric mice

On the day of transplantation, day 10 iPSC-derived human hematopoietic progenitor cells (e/iHPCs) frozen in Cryostor were quickly thawed at 37°C. Cells were then transferred to a 15 mL falcon tube and resuspended by the dropwise addition of 7 mL of DMEM while gently shaking at a 45° angle. Following centrifugation at 500g at 4C, cells were resuspended in 1mL transplantation medium consisting of DPBS (without Ca/Mg) supplemented with 25 ng/mL human M-CSF for counting. Cells were spun at 500g and resuspended in an amount of transplantation media that equals to 500,000 cells in 4 uL transplantation solution. Cells were kept on ice until transplantation.

P2-P4 pups were anesthetized via cryoanesthesia by placing them in a mold on ice until they were unresponsive and lacked a toe-pinch reflex. The pup’s head was then sterilized using alternating swabs of 70% ethanol and betadine. The cell suspension was drawn into a 10 µL Hamilton syringe (7653-01) fitted with a 32-gauge needle (7803-04). Each pup received a subcutaneous injection of Meloxicam (2 mg/kg) for analgesia. Four intracranial injections were performed approximately 2mm along the midline of the skull on both sites equally spaced between lambda and bregma at an approximate depth of 1 mm. Pups were warmed in the experimenter’s hand until fully recovered and ambulatory. Before being returned to their home cage, the pups were rubbed with bedding from the dam to ensure acceptance. Pups were monitored daily for four days post-surgery. Additional doses of Meloxicam (2 mg/kg, s.c.) were administered at 24 and 48 hours post-transplantation to maintain analgesia. Pups were weaned at P21 and aged until tissue collection.

### Tissue collection for brain and leptomeningeal samples

Deep anesthesia of adult mice was achieved by intraperitoneal injection of Avertin (2,2,2-Tribromoethanl in 2-Methyl-2-butanol; Millipore Sigma #T48402 and #240486; diluted in HBSS, ThermoFisher Scientific #14175145), and confirmed using the paw pinch test to verify loss of withdrawal reflex. Then, mice were transcardially perfused with HBSS (≥1mL/g of body weight). HBSS was kept ice-cold for perfusion. Mice were then decapitated and the skull cap was gently removed using forceps and spring scissors to ensure no dural meninges remained attached to the brain and leptomeninges. For immunohistochemical processing, brains (with leptomeninges attached) were then dissected and transferred to 4% PFA (EM grade 32% PFA, Electron Microscopy Science #15714) in HBSS for 24-hour post-fixation. For flow cytometric processing of fresh live tissue, brains (with leptomeninges attached) were carefully dissected and transferred to ice-cold RPMI (RPMI-1640, no phenol red, ThermoFisher Scientific #11835030) containing 10mM HEPES (Sigma-Aldrich #H3537-100; “RPMI-H”) prior to further processing.

### Immunohistochemistry

Paraffin embedding and sectioning was performed at the Boston Children’s Hospital Histopathology Core. Briefly, PFA fixed hemibrains in HBSS were subjected to dehydration steps (50% ethanol, 80% ethanol, 100% ethanol, 100% isopropanol, 50% isopropanol and 50% xylol, twice 100% xylol) followed by embedding sagittally in paraffin. Paraffin-embedded blocks were sectioned sagittally and slices were mounted onto slides and dried overnight and stored at room temperature until staining.

Slides were deparaffinized at 60 °C for 10min followed by incubation in xylol (100% twice) and a 1:1 mixture of xylol and isopropanol for 10 min each. The slides were rehydrated in a descending ethanol series (100%, 90%, 70%, 50%, ddH20) for 5min each. This was followed by incubation in basic antigen retrieval solution (10 mM Tris and 1 mM EDTA pH 10) at room temperature for 5 min and boiling for 10 min. The samples were cooled for 20 min and washed in distilled water for 1 min before subsequent permeabilization in 0.1% Triton X-100 in PBS. This was followed by blocking with 10% HS in PBS for 1 h at room temperature. After blocking, slices were incubated in primary antibody solution (PBS, 10% HS) overnight at 4 °C in coverplates (Epredia). The following antibodies were used in this study: anti-Iba1 (rabbit, Wako; 1:500); anti-human Nuclei (mouse, 3E1.3 Milipor MAB4383; 1:100); anti-mouse CD31 (goat, R&D AF3628; 1:100), anti-human CD163 (mouse, Invitrogen EDHu-1, MA1-82342; 1:500), anti-human Tmem119 (rabbit, Abcam AB185333; 1:100); anti-mouse AQP4 (rabbit, Novus Bio, NBP1-87679; 1:100). Samples were washed for 5min in PBS for three times and incubated with the corresponding fluorescent secondary antibody diluted in PBS containing 10% HS for 2 h at room temperature in the dark. The following fluorescently conjugated secondary antibodies were used: anti-goat Alexa Fluor 755 (donkey, Thermo-Fisher; 1:500); anti-mouse Alexa Fluor plus 647 (donkey, Thermo-FIsher; 1:500); anti-rabbit Alexa Fluor Plus 488 (donkey, Thermo-Fisher; 1:500). Nuclei were stained with Hoechst (1:10,000, Invitrogen H3570) in PBS for 5 min at room temperature. Slides were again washed for 5 min in PBS for three times and mounted with Aqua PolyMount mounting medium (PolySciences), and dried at room temperature.

### Microscopy image acquisition

#### Confocal microscopy

Cover-slipped slides were imaged using Zen Black 2.3 software on a Zeiss LSM880 confocal microscope equipped with 405nm, 458 nm, 488nm, 561 nm, 594 nm, 633 nm, and 730 nm laser excitation. Fluorescent signal was then captured by five detectors: two external (fixed band pass filters; for far-red and infrared signal) and three internal (adjustable band pass filters). Individual z-stacks scans were taken with either a 40x oil-immersion objective. Within confocal microscopy imaging batches, z-resolution, zoom, laser power, detector gain, and aperture were all matched across samples.

#### Slide scanner microscopy

Cover-slipped slides were imaged using a Zeiss Axioscan 7 slide scanner equipped with a Colibri 7 LED source and Zen 3.4 software. Sample detection and section segmentation was done automatically by drawing a marker line around imaging sections prior to imaging. Whole-section images were acquired using a 20x objective and fluorescent imaging using the DAPI, AF488, AF555, AF647, and AF750 channels. Exposure duration and LED power were set per channel. Coarse focus was obtained throughout the section in every tile using the 5x objective and every third row/column strategy on the DAPI channel. Fine focus was obtained for each tile using the onion skin strategy on the AF488 channel. Within slide-scanner imaging batches, all imaging and focus parameters were matched across samples.

### Brain sample preparation for flow cytometric analysis or sorting

#### Brain preparation for acute engulfment assays

Brains were obtained as described in the tissue collection section above. Then, using razor blades, samples were chopped to a fine paste in 200uL of RPMI-H, transferred to FACS tubes (Corning #352058), and mixed into 3mL of ice-cold RPMI-H. Samples were then pelleted via a centrifuge spin (5min x 500g; 4C) to enable removal of the supernatant and thorough re-suspension in digestion buffer (2mL for half brains, 4mL for whole brains). The digestion buffer contained 0.5 mg/mL Collagenase P (Sigma #11213865001), 0.8 mg/mL Dispase II (Worthington #LS02104), 250 U/mL DNAse-1 (Worthington #LK003172), and 1µM pre-formed Aβ_1-42_ fibrils (StressMarq Biosciences #SPR-487E) in RPMI-H, and was pre-warmed at 37C prior to use. Prior to inclusion in the digest buffer, synthetic Aβ_1-42_ fibrils were conjugated to pHrodo Deep-Red (Invitrogen #P35358) according to manufacturer instructions. Samples were light-protected, incubated at 37C, and gently agitated using an orbital rocker for the 30 minutes of enzymatic digestion.

#### Brain preparation for FACS prior to single-cell RNA sequencing

Brains were obtained as described in the tissue collection section above with the exception that transcardial perfusion was performed using a buffer containing transcriptional and translational inhibitors to prevent *ex vivo* transcriptional artifact^23^: Actinomycin D (5µg/mL, Sigma-Aldrich #A1410) and Triptolide (10µM, Sigma-Aldrich #T3652) in RPMI-H. Samples were then incubated and minced in a solution of 5µg/mL Actinomycin D, 10µM Triptolide, and 27.1µg/mL Anisomycin (Sigma-Aldrich #A9789) in RMPI-H, and digested for 30 minutes at 37C with gentle agitation in a digest buffer containing 0.5 mg/mL Collagenase P, 0.8 mg/mL Dispase II, 250U/mL DNAse-1, 5µg/mL Actinomycin D, 10µM Triptolide, and 27.1µg/mL Anisomycin in RPMI-H. All solutions and samples were kept light-protected.

#### Cell purification and labeling for flow cytometry

Following digestion, all steps prior to flow cytometric analysis were performed on ice or at 4C and using ice-cold reagents to prevent artifact. First, samples were spun to pellet the tissue (5 minutes x 500g). The supernatant was removed and tissue pellets were resuspended and gently triturated in 1mL of FACS buffer, composed of 0.5% Bovine Serum Albumin (BSA, Sigma-Aldrich #A2153) and 2mM EDTA (Research Products International #E14000) in HBSS. Following trituration, samples were filtered through a 70µm filter and diluted in 25% BSA in HBSS to a final volume of 7mL for a cellular enrichment and debris removal spin (10 minutes x 1200g). Careful to avoid disrupting the cell pellet, the resulting myelin layer and supernatant were removed. Then, the cell pellet was re-suspended in 1mL of FACS buffer, transferred to a 2mL low-bind tube, and centrifuged again (5 minutes x 500g). Following supernatant removal, the cell pellet was re-suspended in 200uL of FACS buffer and transferred to 96-well U-bottom plates (Thermo Scientific #262162). Cells were pelleted once more using the centrifuge (5 minutes x 500g), then resuspended in 50µL of an Fc-blocking and viability dye solution containing 2% Fc-blocking antibody (anti-mouse CD16/CD32, BD Biosciences #553141) and 0.1% viability dye (eBioscience Fixable Viability Dye eFluor 780, ThermoFisher Scientific #65-0865-18) in FACS buffer. After 10 minutes of incubation, 50µL of antibody cocktail solution were added into each well and gently pipette-mixed, ensuring to avoid bubbles. Antibody cocktails were made in FACS buffer containing 20% Brilliant Stain Buffer (BD #566385), and all antibody clones, catalog numbers, and dilutions are listed in **Table S3**). Samples were stained for 20 minutes on ice, then washed twice in 200uL FACS buffer with 5 minutes x 500g centrifuge spins prior to flow cytometric analysis or sorting.

#### Sample preparation and staining for fixed-cell FEAST protocol

We used FEAST to assess *in vivo* MBP engulfment in control and cuprizone-demyelinated mice^32^. Importantly, the fixed-cell protocol was used as live cell preparations were shown to incompletely prevent *ex vivo* engulfment by BAMs^32^. All steps were performed as described in the protocol. Briefly, mice were transcardially perfused with 10mL of HBSS and then 20mL of fixation buffer (BioLegend #420801). Brain and leptomeningeal samples were dissected as described above and then incubated in a fixation quenching solution containing 5mM HEPES (Sigma-Aldrich #7365-45-9), 250mM Tris (Millipore Sigma #GE17-1321-01), and 250mM Glycine (Millipore Sigma #G7126) in HBSS. Samples were minced to a fine paste using a razor blade and underwent a 2-hour enzymatic digestion at 37C in 800U/mL of Collagenase IV (Worthington #LS004189) in RPMI-H with gentle agitation. Then, samples were triturated and processed using a BSA enrichment spin, an acid wash, and a blocking step as described in the fixed cell FEAST protocol to yield single-cell suspensions while minimizing debris and contamination from material adhered to the cell surface^32^. Prior to staining, single cell suspensions underwent a second fixation in IC fixation buffer (eBioscience #00-8222-49), and were then blocked and permeabilized permeabilization buffer (eBioscience #00-8333-56) containing 2% Fc-blocking antibody (anti-CD16/CD32, BD Bioscience). Samples were stained using antibodies against both human antigens (CD45, CD206), murine antigens (CD45, GR-1, CD68, CX3CR1, CD38, MHC-II), and against MBP, in addition to 1:20,000 DAPI (BioLegend #422801). All antibody specifications including clone number, catalog number, and recommended dilution are listed in **Table S3**.

### Flow cytometric analysis and sorting

Flow cytometric analysis and fluorescence-activated cell sorting (FACS) were performed using a FACSSymphony S6 flow cytometer using FACSDIVA v9.5 software. For analysis, stained single-cell suspensions in FACS buffer were transferred to FACS tubes through a 35µm filter cap prior to loading into the machine. All samples within a given experiment were processed jointly in a single analysis session, and FSC files were subsequently exported for analysis using FlowJo v10.10.0. For FACS, single-cell suspensions in a coating buffer containing just HBSS and 2% BSA were transferred to FACS tubes through a 35µm filter cap prior to loading into the machine. Cells were sorted into 2mL low-bind microcentrifuge tubes containing coating buffer and using the gating strategy shown in **Figure S2A**.

### Single-cell RNA sequencing

#### Library preparation and sequencing

Following FACS, cells were pelleted at 4C using a centrigue (5 minutes x 500g) and the supernatant was removed, leaving ∼30μl. The cell pellet was then gently resuspended using a pulse vortex and micropipettor to a final volume of 38.7μl. Then, droplet generation was performed using the 10X Genomics 5’ Single-cell Version 2 and NextGEM Chromium Controller and barcoded libraries were generated as per manufacturer specifications. Then, single-cell libraries underwent an initial shallow-depth sequencing step using the Illumina NextSeq 500 with a High Output 150 cycle flowcell. For this step, single-cell libraries were pooled at equimolar concentrations prior to denaturation and dilution following Illumina guidelines for NextSeq500 High Output flow cells. Subsequently, single-cell libraries underwent full-depth sequencing. For this, libraries were re-pooled based on parameters established from shallow depth sequencing, notably cell number, to match read depth per cell across all libraries. The pooled library was then sequenced on a NovaseqX Plus 10B flowcell at the Broad Institute’s Genomics Platform.

#### Single-cell Data Preprocessing

The following preprocessing was performed on the Broad Institute’s Google Cloud Workspace. Single-cell RNA sequencing data was aligned to the combined human GRCh38 and mouse GRCm39 genome (release 2024) using 10x Genomics Cell Ranger (v9.0.2) following standard Cell Ranger count workflow and ‘--create-bam true’. Ambient RNA was subsequently removed using ‘remove-background’ from CellBender v0.3.0 (Fleming et al. 2023). The following settings were used for CellBender processing: ‘expected_cells’ was set equal to Cell Ranger called cells, ‘total_droplets_included’ set based on examination of barcode rank plots, ‘--epochs=150’ and ‘--learning-rate=1e-4’.

#### Quality control

All further processing and analysis was conducted using Seurat v5. A mitochondrial score for human and murine cells was calculated using the gene prefixes of “GRCh38-MT-” and “GRCm39-Mt-”, respectively. Then, the dataset was filtered to remove cells with <500 detected genes or cells with either 0% or >10% of transcripts mapping to human or murine mitochondrial genes. The filtered object was then processed following the standard Seurat v5 workflow^50,51^. Briefly, the object underwent log-normalization, variable feature detection, data scaling with regression to RNA count and mitochondrial score, determination of dimensionality, and clustering. In this initial clustered object, individual cell types were then subsetted for quality control (QC) using a sequential high-resolution clustering approach to identify outlier clusters. Outlier clusters of low quality cells to exclude were identified based on lack of transcripts mapping to ribosomal genes or an aberrantly high proportion of transcripts mapping to mitochondrial genes combined with a low feature RNA number. We also used this approach to identify doublets and multiplets. Clusters containing aberrantly high feature and RNA count in addition to incongruent cell type-specific markers were considered multiplets and excluded (evident multiplets, e.g. a BAM cluster enriched for endothelial cell-specific genes, or a cell containing both human and murine RNA). Each round of high-resolution clustering was sequentially repeated until all low-quality cells and multiplets were identified and removed. All cells that passed this QC pipeline were subsetted from the original Seurat object into a final object for analysis. This final object underwent normalization, feature detection, scaling, and dimension reduction prior to analysis.

#### Annotation and data visualization

We performed both a broad and fine annotation. For both rounds of annotation, we used the Seurat v5 FindAllMarkers function with the following parameters: min.pct = 0.25, logfc.threshold = 0.405. For broad annotation, FindAllMarkers was performed on the whole object. For fine annotation, cells were subsetted based on species origin, and FindAllMarkers was performed on each subset. For visualization of feature expression, clusters, and metadata, we used uniform manifold approximation and projection (UMAPs) and overlaid feature plots. Complex heat maps and sacked violin plots were made using scCustomize (version 3.0.1)^52^. To generate replicate-level heatmaps, we used the AggregateExpression function in Seurat v5 (grouped by orig.ident and cell type) paired with the DoHeatmap function using scaled data. K-means clustering of BAMs and microglia across Zeitgeber Time using the conserved gene set was performed using the Clustered_DotPlot function of scCustomize (version 3.0.1).

#### Identification of the conserved BAM versus microglia gene set

We first performed differential gene expression analysis to obtain markers for BAMs and xBAMs, separately. For this, we used two subsetted objects: one of just BAMs and microglia, another of just xBAMs and xMGs. On each object, we performed cell-level differential gene expression analysis using the FindAllMarkers function with the following parameters: only.pos set to FALSE, min.pct set to 0.4, and logfc.threshold set to 1, using the “MAST” package for differential expression testing^53^. The output was filtered to only contain genes enriched or de-enriched in BAMs/xBAMs with an adjusted p-value <0.05.

The convert_orthologs() function available from the orthogene (v1.14.0) Bioconductor package was then used to determine conserved genes between the murine and human output. Two rounds of ortholog verification were performed: 1) identifying mouse orthologs from the human DEG output, and 2) identifying human orthologs from the murine DEG output. For each set of inputs, genes from both species were dropped if there was not a 1:1 gene mapping. Orthologs were identified using the gProfiler method within orthogene. Custom R code was used to identify conserved genes between the two orthogene outputs, and to determine which genes were enriched in the same or different directions. The final conserved BAM versus microglia gene set only contained genes that were: 1) identified as orthologs via the two-step verification, and 2) enriched in the same direction in BAMs and xBAMs (**Table S2**). To determine cellular pathways enriched in the BAM signature, we performed pathway enrichment analysis using Ingenuity Pathway Analysis^54^ (Qiagen) on the gene set from the conserved signature with a logFC>1.

### Bulk RNA sequencing of iMGLs

#### Collection and RNA isolation

iMGLs plated in matrigel-coated 24-well plates were used for RNA isolation and bulk-sequencing analysis. On the day of harvest (day 44) media and floating cells were fully removed and adherent cells were immediately lysed in 350uL RLT buffer (Qiagen) supplemented with 40mM DTT. RNA was isolated RNeasy Micro Kit (50) (Qiagen #74004) following manufacturer’s standard protocol. Bulk-RNA sequencing was performed using Novogene services (non-direction, NovaSeq X Plus Series (PE150) generating 6 Graw data per sample). Novegene performed RNA sample quality control as well as mRNA library preparation (poly A enrichment) and sequencing.

#### Bulk RNAseq alignment and quality control

Raw sequencing reads (FASTQ files) were processed to quantify transcript abundances. Salmon (v1.10.3) was used (cite https://www.nature.com/articles/nmeth.4197) in quasi-mapping mode to quantify expression against an index built from the human transcriptome (GENCODE, corresponding to genome assembly GRCh38.p14). The mapping-based mode was run with Salmon standard parameters. Further analysis was performed using Python (v3.12.9) utilizing pytximport package^55^ (v0.12.0) to summarize transcript counts to the gene level and create a gene-by-sample count matrix for downstream differential expression analysis. Initial quality control was performed on the gene count matrix.

#### Bulk RNAseq differential gene expression analysis

Differential gene expression analysis was performed using the pydeseq2 package^56^ (v0.5.0). Raw gene counts were used as input for a DeseqDataSet object. A principal component analysis (PCA) was conducted on variance-stabilized expression data to visualize sample-to-sample distances and identify outliers. Based on this analysis, all samples clustered with their respective biological replicates and were retained for downstream analysis. The analysis followed the standard workflow including size factor estimation for library size normalization, gene-wise dispersion estimation, and fitting a negative binomial generalized linear model to identify differentially expressed genes between experimental conditions. The significance of expression differences was determined using the Wald test, and the resulting p-values were adjusted for multiple testing using the Benjamini-Hochberg method. Genes with padj < 0.05 were considered significantly differentially expressed. For annotation, ENSEMBL gene IDs were converted to official gene symbols using the sanbomics tools.map function (v0.1.0). To identify over-represented biological themes, Gene Ontology (GO) enrichment analysis was performed. This analysis was conducted using the gprofiler-official Python package (v1.0.0) with the human GO:BP 2025 library term.

#### Bulk RNAseq data visualization

To visualize the results, heatmaps of differentially expressed genes were generated using the PyComplexHeatmap library (v1.8.2) (cite https://onlinelibrary.wiley.com/doi/10.1002/imt2.115). Expression values were transformed into Z-scores for each gene across samples to allow for relative comparison, and both genes and samples were clustered using hierarchical clustering. Volcano plots were created to visualize the relationship between the magnitude of gene expression change (log2 fold change) and statistical significance (adjusted p-value). To compare differentially expressed gene lists between different experimental conditions, Venn diagrams were generated using the pyvenn package (v0.1.3). Unless stated otherwise, the analysis and visualization pipeline was conducted using the pandas (v2.2.3), numpy (v1.26.4), and matplotlib (v3.10.1) and scipy (1.15.2) libraries.

### Statistical analyses

Statistical analyses were performed using R (version 4.4.2) driving RStudio (version 2024.09.1+394, Posit PBC) or JMP Pro (version 18, JMP Statistical Discovery). Statistical testing on scRNAseq data is described in the single-cell RNA sequencing methods section. For other comparisons, the statistical test used is indicated in figure legends. An alpha cut-off of 0.05 was used unless otherwise specified. For comparisons between two independent groups, we used either parametric or non-parametric comparisons depending on data distribution and sample size: two-tailed unpaired Student’s t-tests or Mann-Whitney U (Wilcoxon rank-sum) tests, respectively. Grouped comparisons were performed using one- or two-way ANOVAs. In some instances, mixed effects models using restricted maximum likelihood (REML) approaches were used to account for random variables (for example, experimental batch), in which case model parameters are included in the figure legend. For all ANOVAs and mixed effects testing, main and interaction effects are indicated and post-hoc testing was performed with Tukey’s HSD (Honestly Significant Difference). Data visualization was performed using Rstudio, JMP Pro, and FlowJo.

## RESOURCE AVAILABILITY

All sequencing data will be deposited in the Gene Expression Omnibus (GEO) upon manuscript publication. No new reagents were created within the scope of this study, however, requests for protocol sharing are gladly accepted.

## ACKNOWLEDGEMENTS

We thank Liza Curtis, Krishna Narayanan, and Anna Kane for administrative support and Ronald Mathieu from the HSCI-BCH Flow Cytometry Research Laboratory for technical expertise. This work was generously supported by grants from the Simons Foundation Collaboration on Plasticity and the Aging Brain (B.S.), the Cure Alzheimer’s Fund (B.S.), a NIH Training Grant on Molecular Biology of Neurodegeneration and Alzheimer’s Disease T32AG000222 (H.J.B.), and the Howard Hughes Medical Institution (B.S.). BioRender was used for schematic illustrations.

## AUTHOR CONTRIBUTIONS

Conceptualization, H.J.B. and C.M.D.; Methodology, H.J.B., C.M.D., M.H., T.N., S.E.M.; Investigation, H.J.B., C.M.D., M.H., T.N., J.P., S.E.M.; Software, H.J.B., M.H., J.M.R., S.E.M., Supervision, H.J.B., C.M.D. and B.S.; Formal analysis, H.J.B., C.M.D., M.H., J.M.R.; Visualization, H.J.B., C.M.D., M.H.; Project administration, H.J.B. and C.M.D.; Writing – original draft, H.J.B., C.M.D., M.H., B.S.; Funding acquisition, H.J.B., C.M.D., B.S.

## SUPPLEMENTAL INFORMATION

Document S1: Figures S1-3 and Tables S1-4.

## SUPPLEMENTAL FIGURES

**Figure S1.**
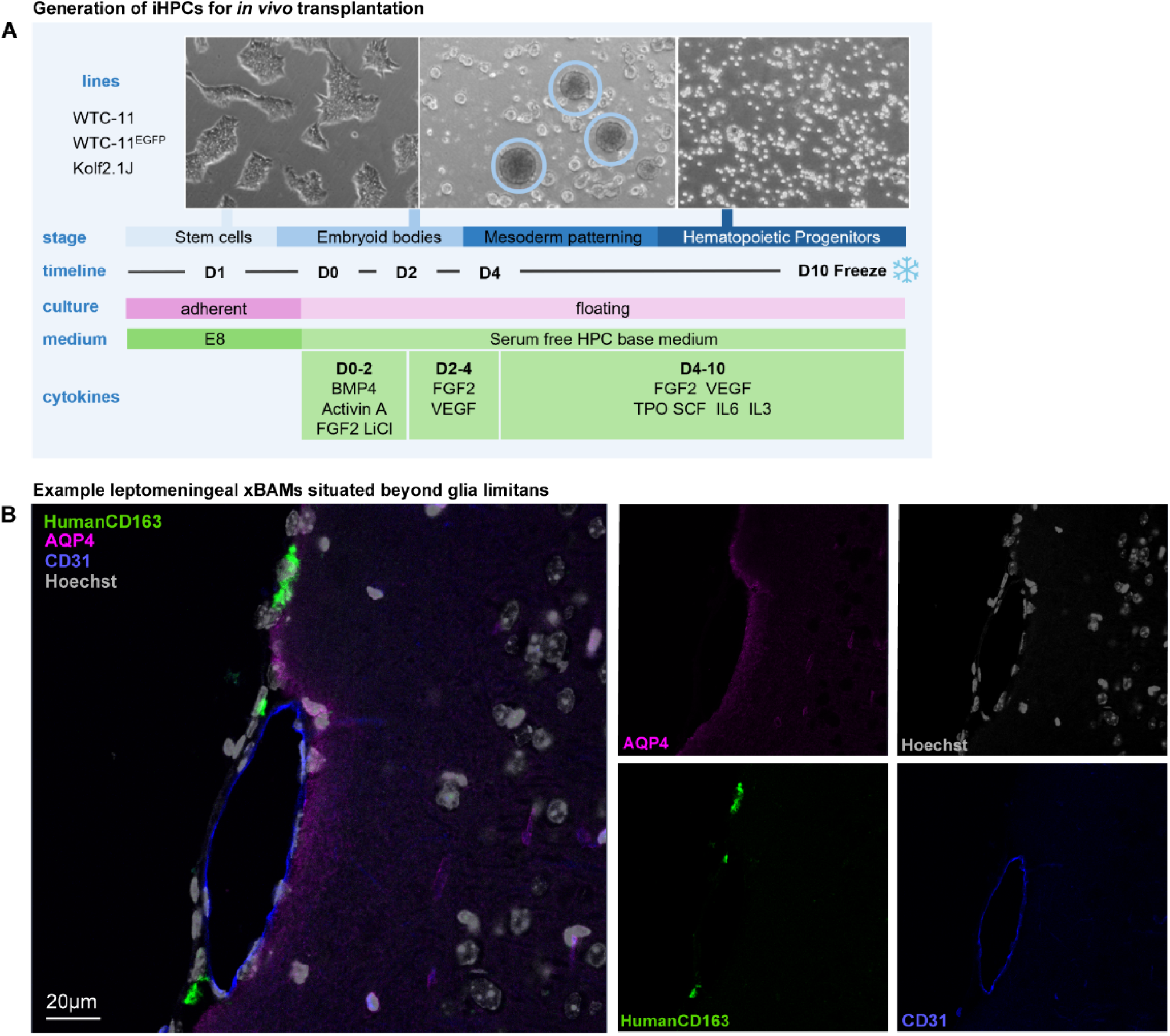
Further characterization of chimeric human-murine brain immune compartment. (**A**) Schematic diagram of experimental design for generating iPS-derived HPCs for transplantation (see **Methods**). (**B**) Example image of human xBAM (labeled with anti-human CD163) located along a pial vessel (labeled with anti-mouse CD31) in the leptomeninges (delineated with anti-mouse AQP4).

**Figure S2.**
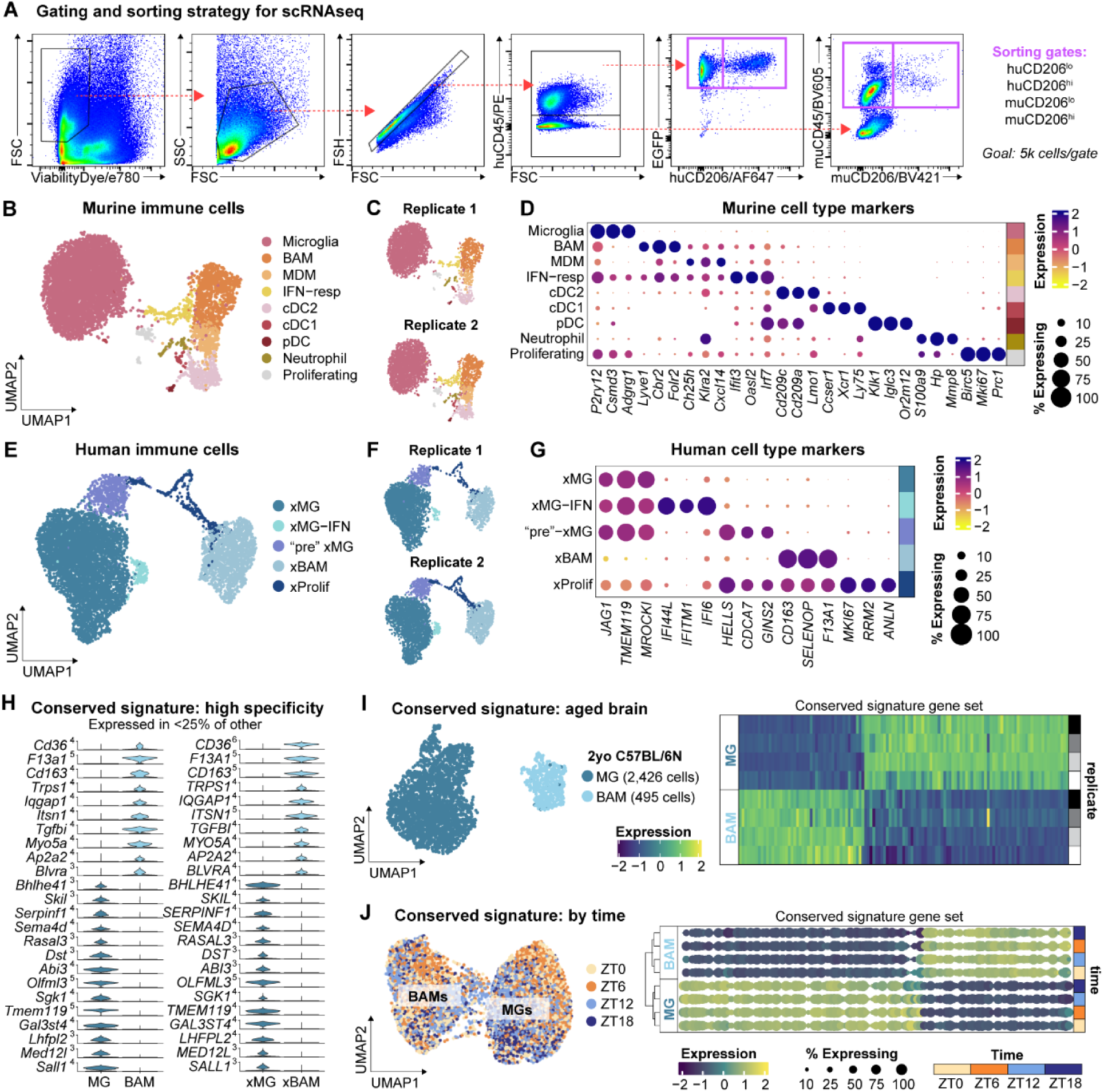
Further characterization of conserved BAM versus microglia signature. (**A**) Flow cytometry gating and sorting strategy for scRNAseq of chimeric immune compartment. Purple boxes represent sorting gates. (**C-D**) UMAP of annotated murine immune cell subsets, UMAP split by biological replicate, and heat map of murine immune cell type markers. cDC1/2 = conventional dendritic cell type 1/2, pDC = plasmacytoid DC. (**E-G**) UMAP of annotated human immune cell subsets, UMAP split by biological replicate, and heat map of human immune cell type markers. (**H**) Violin plot of highest specificity BAM vs. microglia markers, indicated by conserved expression under 25% in the opposing cell type across both human and murine cells. (**I**) UMAP of microglia and BAMs from 2 year-old C57BL/6 male mice (n of four biological replicates) and heatmap of expression of conserved signature gene set across cell types. (**J**) UMAP of microglia and BAMs from 2.5 month-old C57BL/6 male mice (n of 12 biological replicates total, 3 replicates per time point) grouped by time and k-means clustering of cell types by time using conserved signature gene set. Regardless of time of day, hierarchical clustering still distinguishes BAMs and microglia using only the conserved signature gene set.

**Figure S3.**
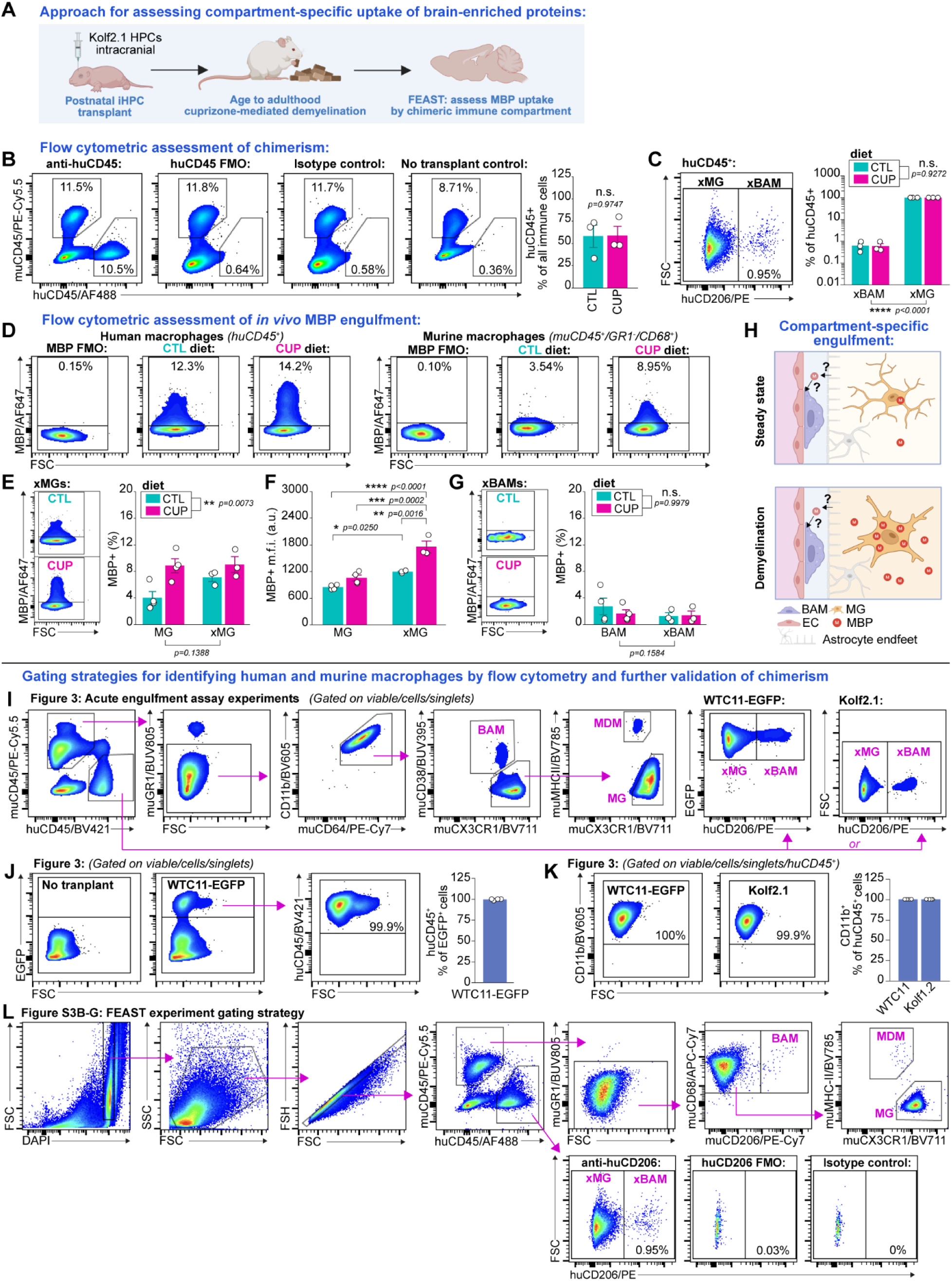
A flow cytometric toolkit for assessing *in vivo* function of human brain macrophages. (**A**) Schematic diagram of experimental design. Mice received a postnatal transplant of iPS-derived HPCs (Kolf2.1). In adulthood, mice underwent cuprizone-mediated demyelination and MBP uptake was assessed across the chimeric immune compartment via FEAST. (**B**) Pseudo-colored plots demonstrating specificity of anti-human CD45 staining in chimeric brains and group-level quantification of chimerism across conditions. CTL = control diet, CUP = cuprizone diet. No effect of diet on chimerism (two-tailed student’s t-test; n of 3 chimeras per diet). (**C**) Pseudo-colored plots demonstrating specificity of anti-human CD206 staining in chimeric brains and group-level quantification of distribution of chimeric cell types across conditions. No effect of diet on composition of chimeric immune compartment (two-way ANOVA; main effects shown; diet*cell type interaction effect: p=0.8226; n of 3 chimeras per diet). (**D**) Representative pseudo-colored plots of MBP uptake by macrophages of human and murine origin across control and cuprizone diets, with negative control MBP staining. (**E and F**) Representative pseudo-colored plots of MBP signal in xMGs from control and cuprizone-treated mice, and group-level quantification of MBP-positivity and signal in MGs and xMGs across conditions. Main effect of diet but not cell type, and no interaction between the two (p=0.1813), on MBP-positivity, and cell type*diet interaction effect on MBP signal within MBP+ cells: p=0.0315. Quantifications using two-way ANOVAs, post-hoc testing with Tukey’s HSD, n of 3-4 per condition. (**G**) Representative pseudo-colored plots of MBP signal in xBAMs from control and cuprizone-treated mice. No effect of diet or cell type on MBP uptake by BAMs and xBAMs (two-way ANOVA; main effects shown; diet*cell type interaction effect: p=0.6086; n of 3-4 per condition). (**H**) Schematic diagram of compartment-specific engulfment. Cuprizone-mediated demyelination induces robust increases in MBP uptake by MGs and xMGs but not BAMs/xBAMs. (**I**) Gating strategy for acute *ex vivo* engulfment assays in Figure 3. (**J**) Verification of fate of xenotransplanted human cells. Representative pseudo-colored plots of EGFP expression in WTC11-EGFP-transplanted mice, and huCD45 expression within EGFP^+^ cells. The proportion of EGFP^+^ cells that are huCD45^+^ is not different from 100% (two-tailed one-sample t-test against mean of 100, p = 0.3163; n of 4 independent replicates). All xenotransplanted cells detected by flow cytometry had thus become CD45^+^ leukocytes. (**K**) Verification of identity of xenotransplanted human cells. Representative pseudo-colored plots of CD11b expression in huCD45^+^ cells from WTC11-EGFP- and Kolf2.1-transplanted mice. 99.9±0.01% of all huCD45^+^ were CD11b^+^ (mean±SEM, n of 4 per condition). All xenotransplanted leukocytes detected by flow cytometry had thus become CD11b^+^ macrophages. (**L**) Gating strategy for FEAST experiment in Figure S3B-G. *For all panels: points represent individual mice, bars and error bars represent mean and SEM. Illustrations made with BioRender*.

## SUPPLEMENTAL TABLES

**Table S1.**
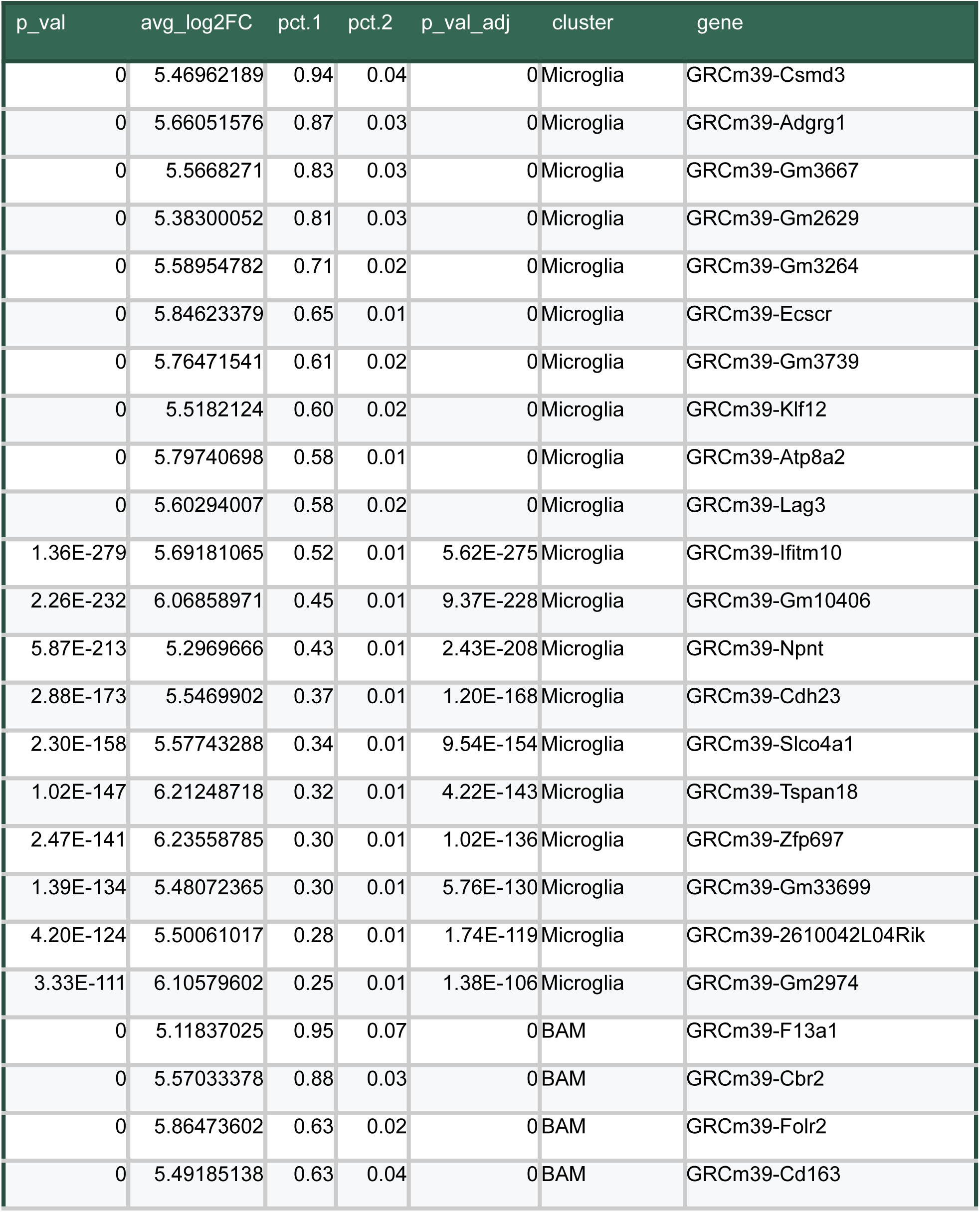
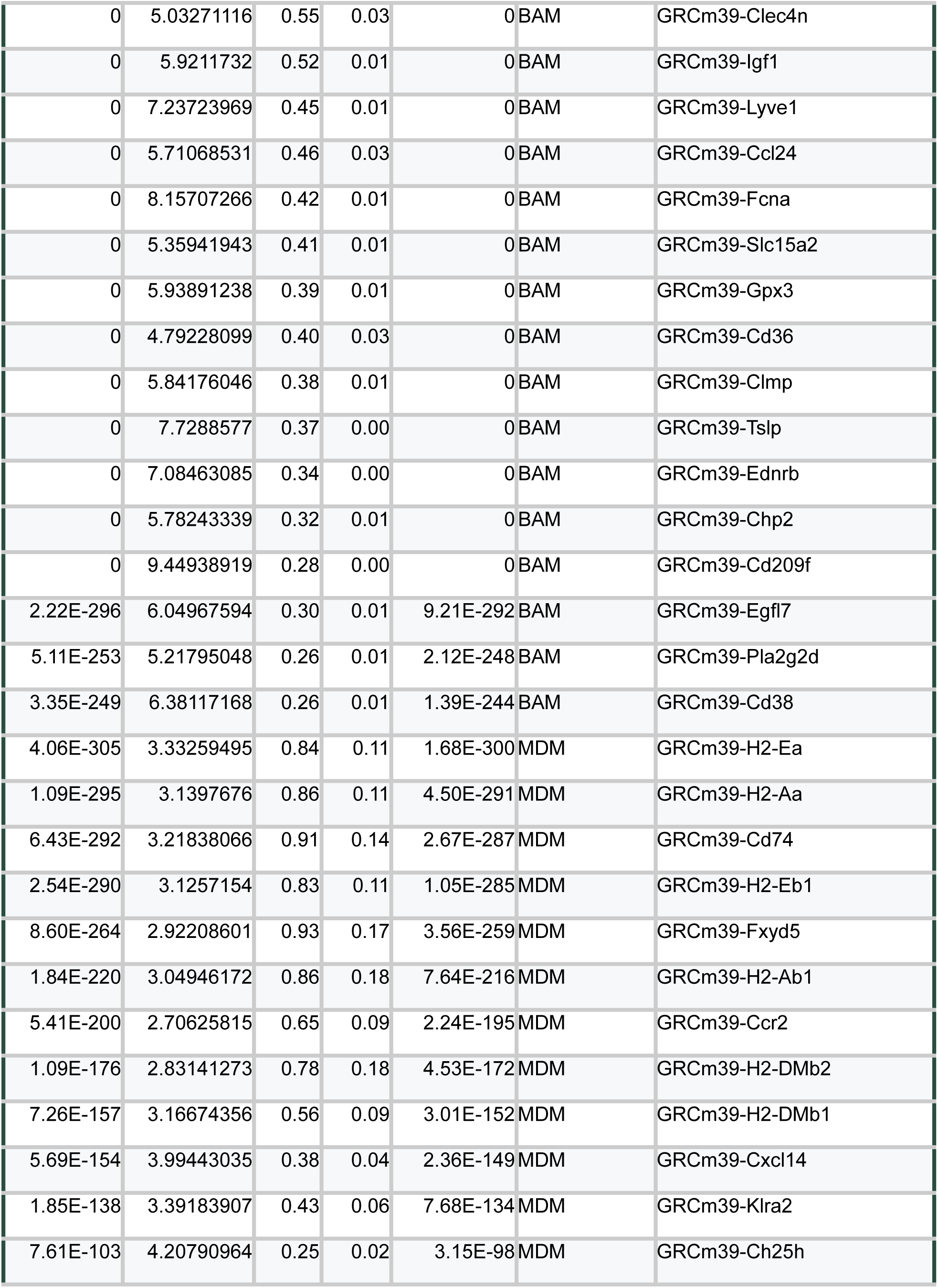
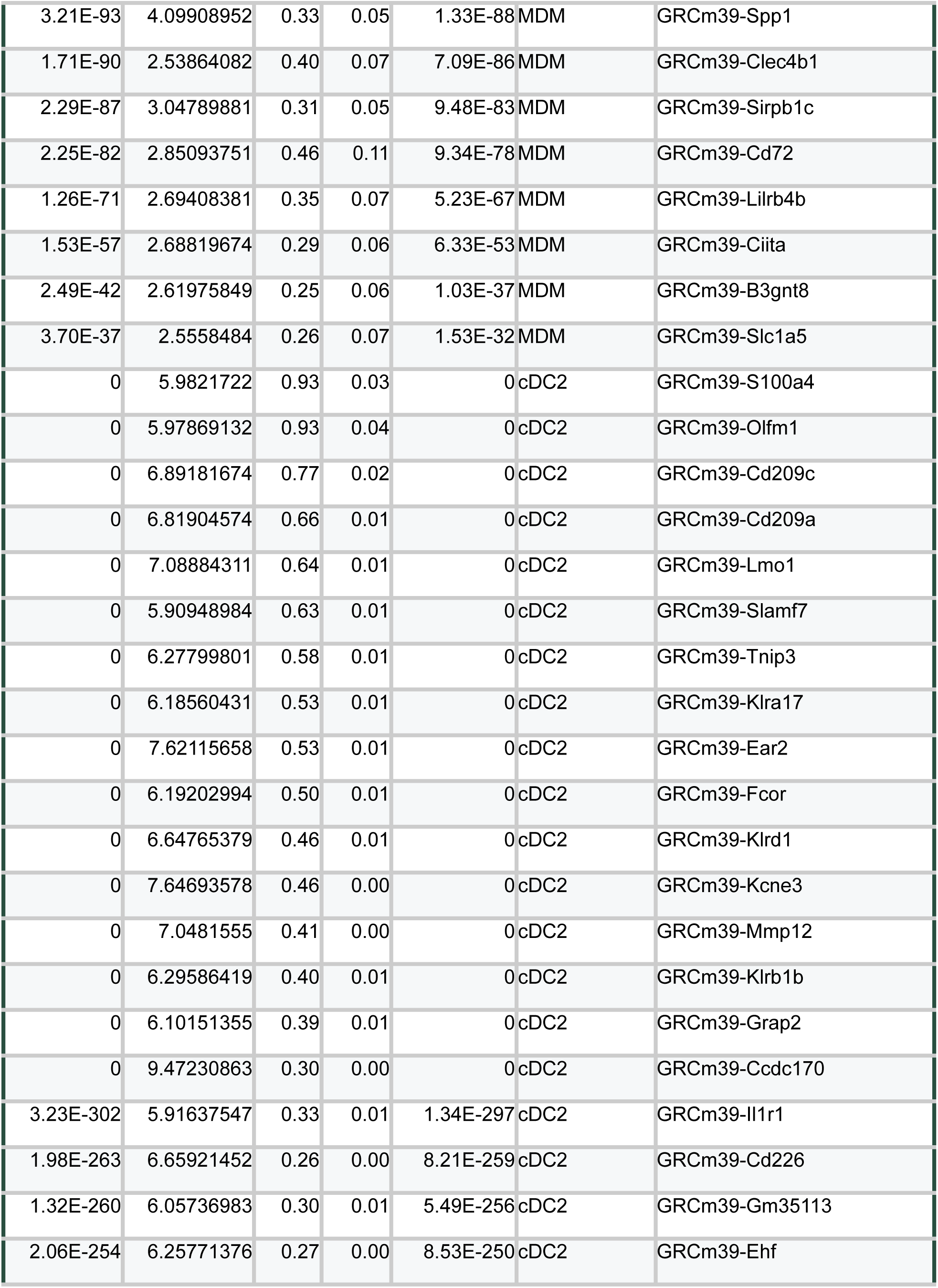
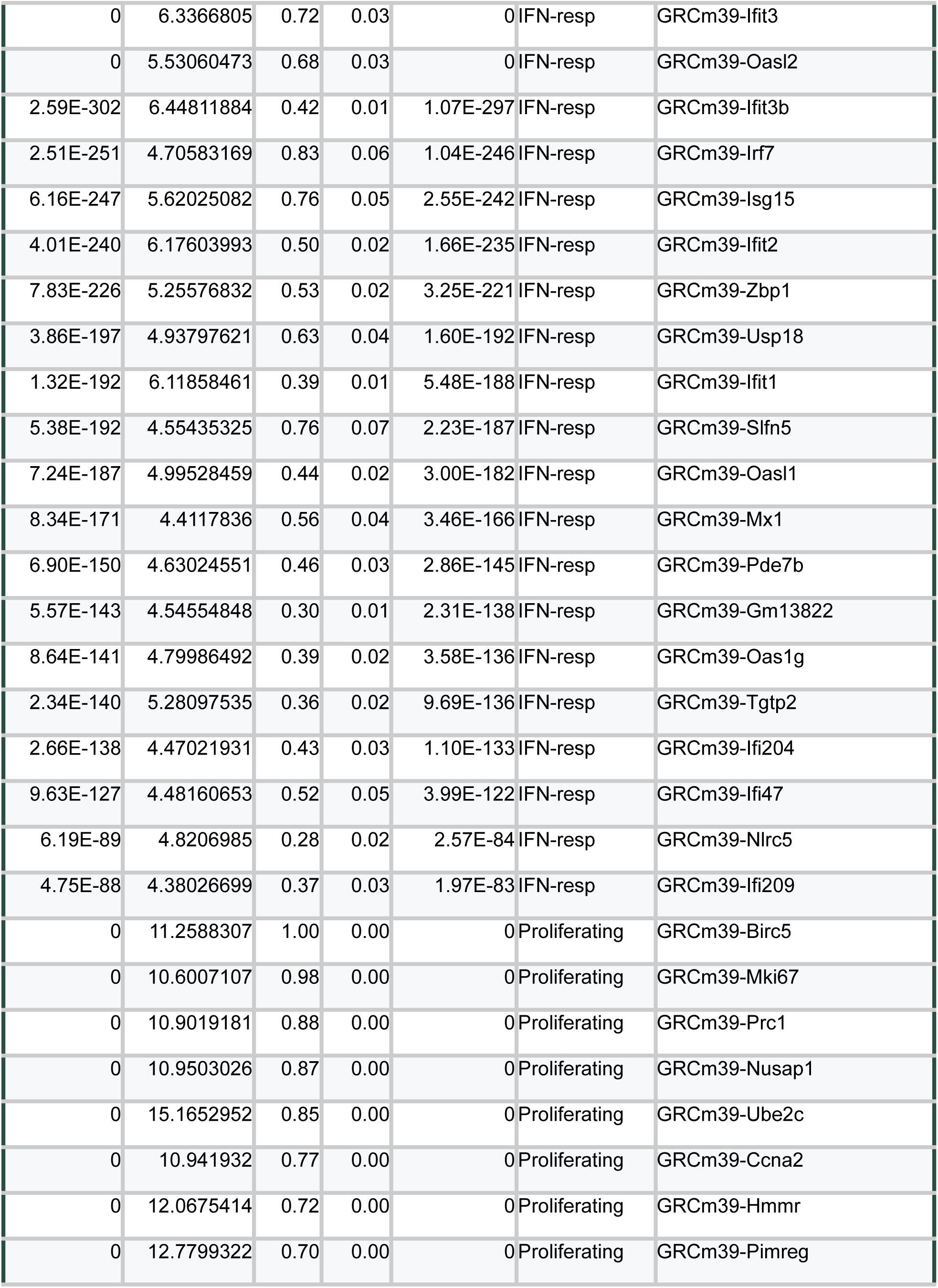
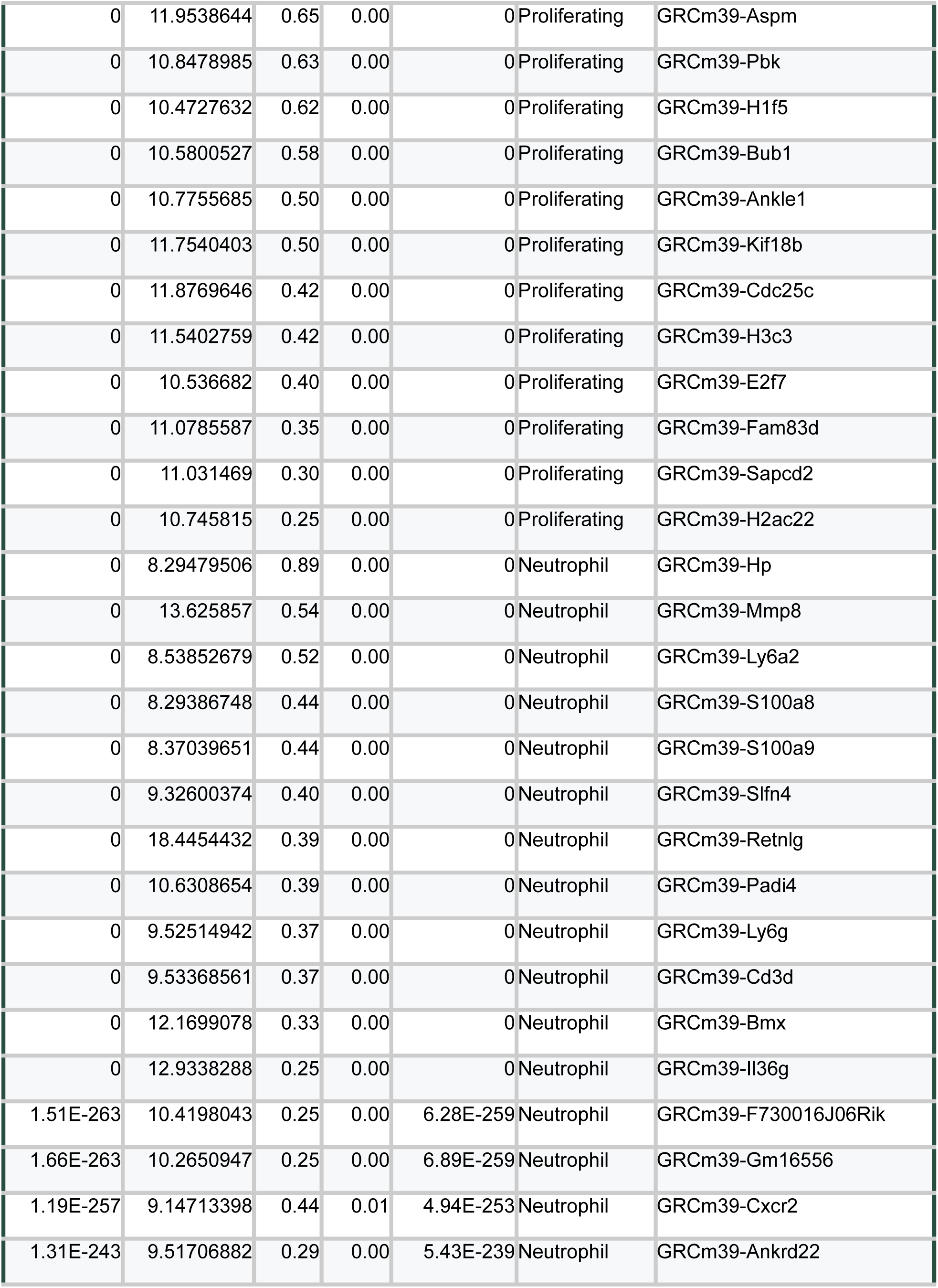
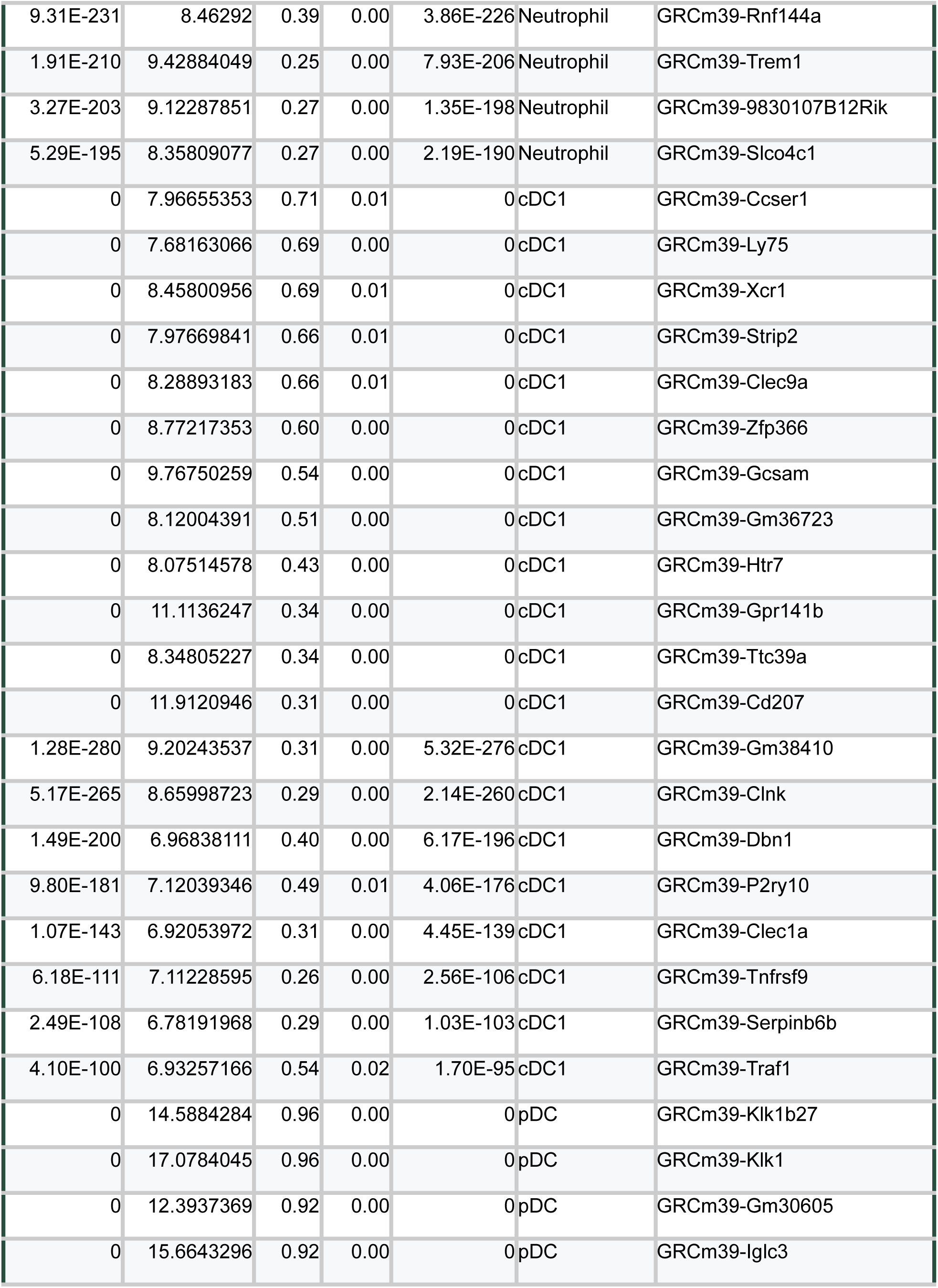
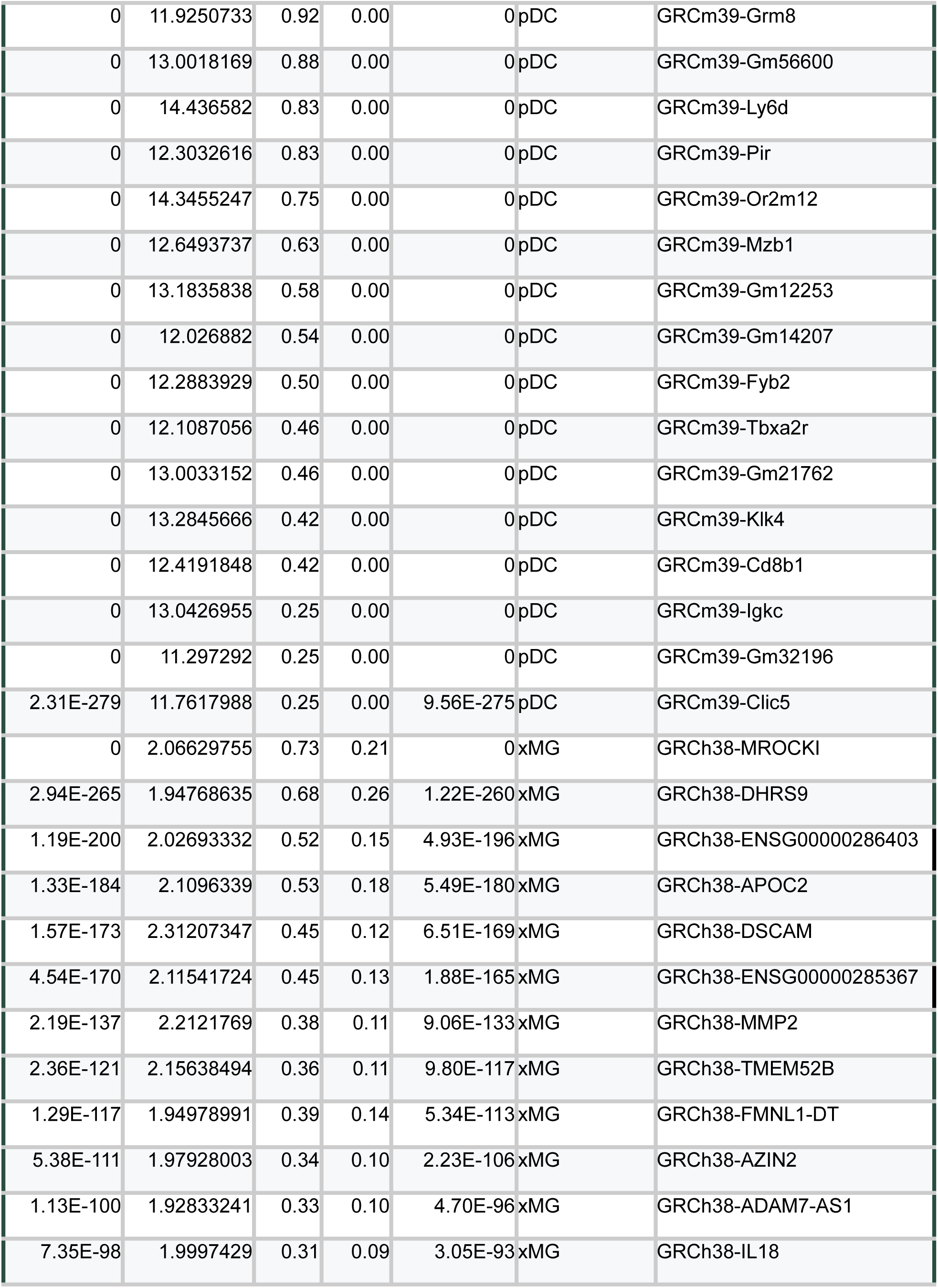
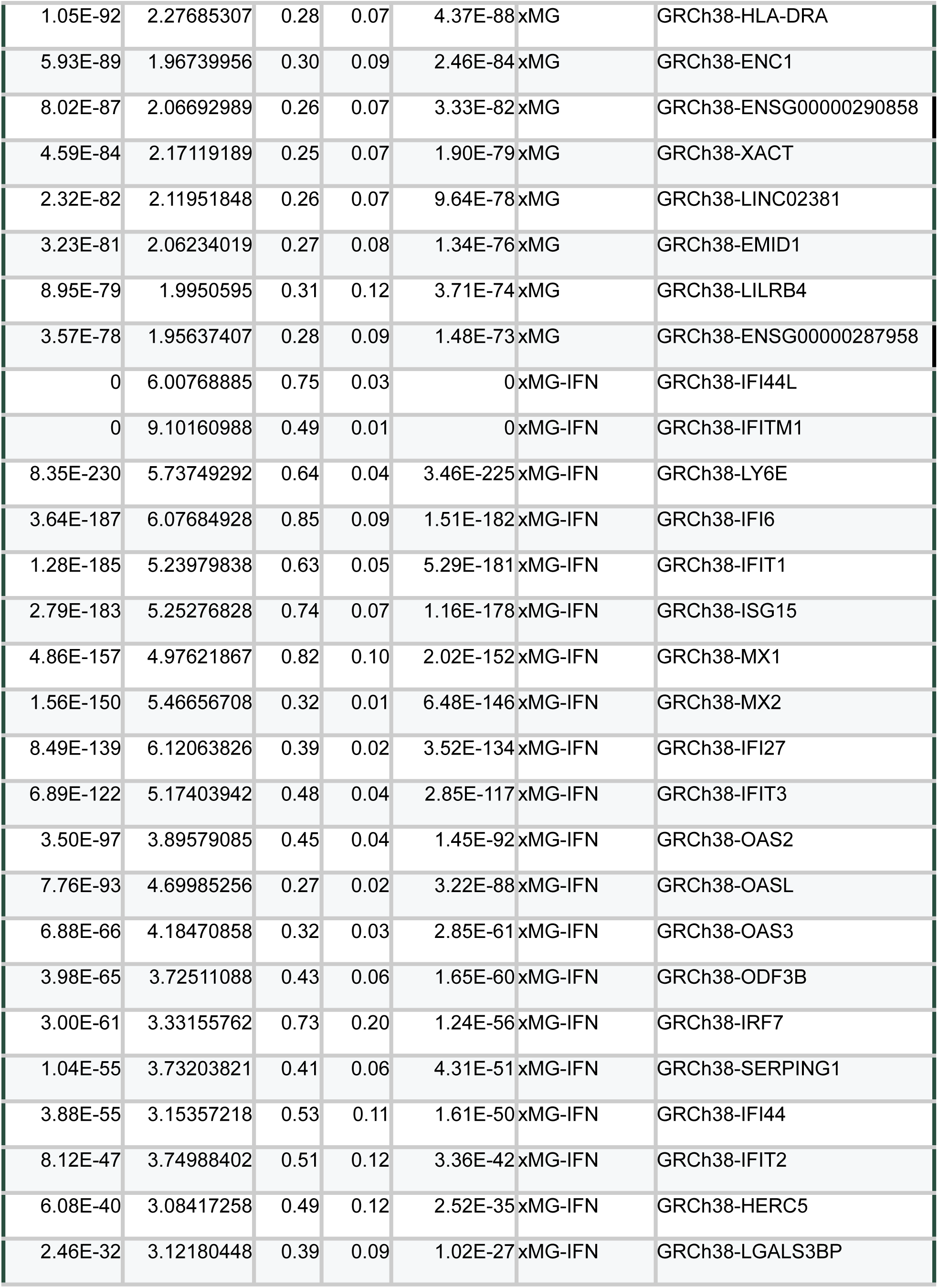
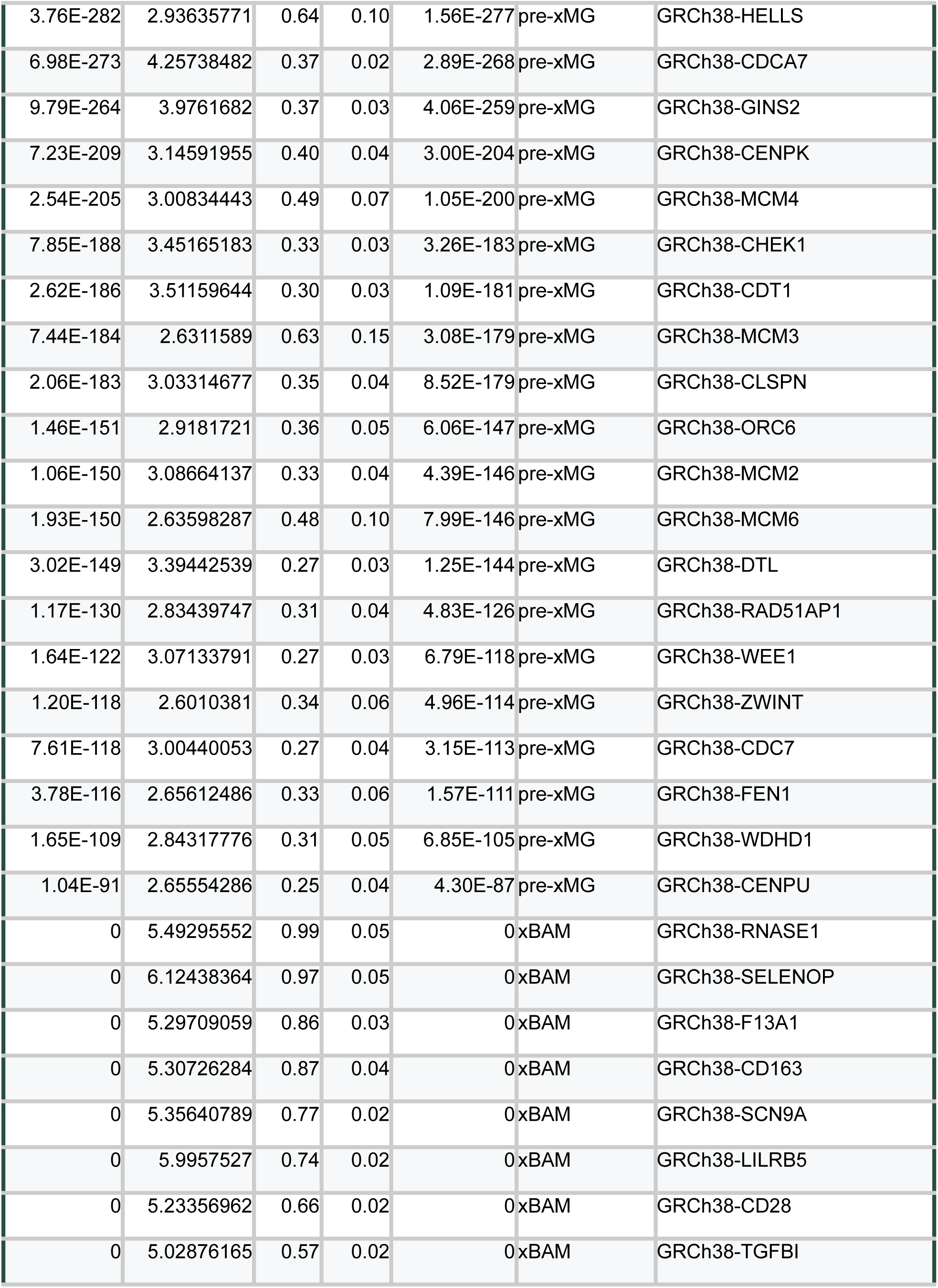
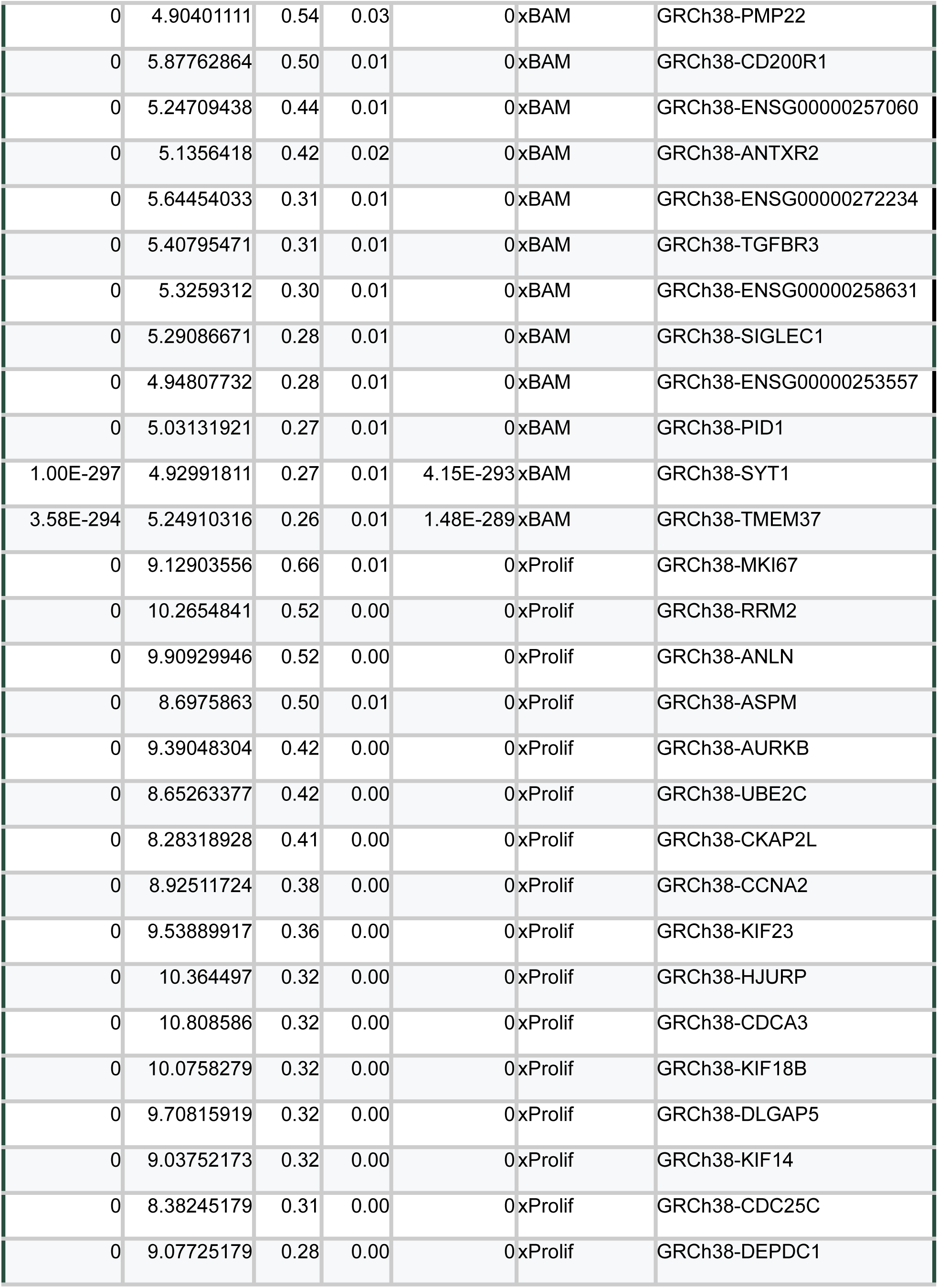
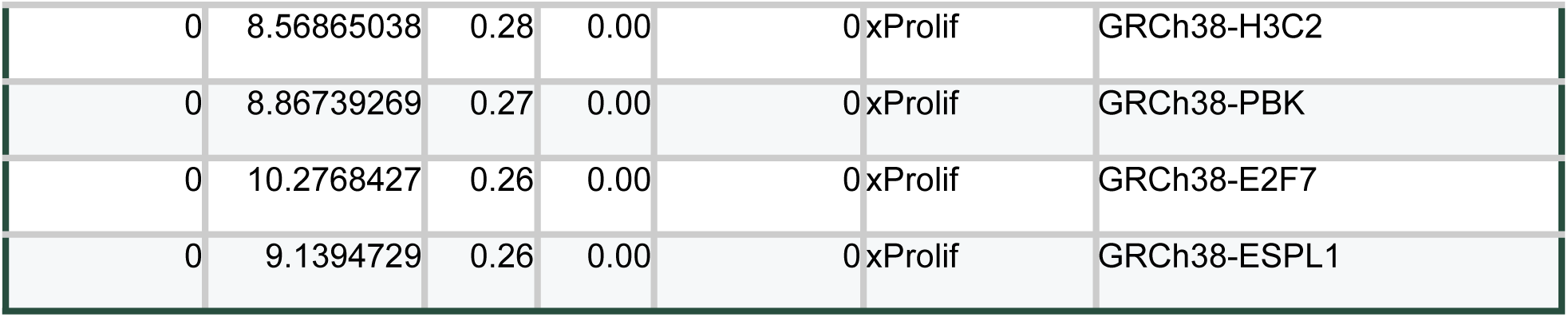
Top genes per cell type in scRNAseq dataset. Marker genes per cell type. Prefix in gene name corresponds to genome annotation. GRC = Genome Reference Consortium, GRCm39 = Mus musculus 39, GRCh38 = Homo sapiens 38.

**Table S2.**
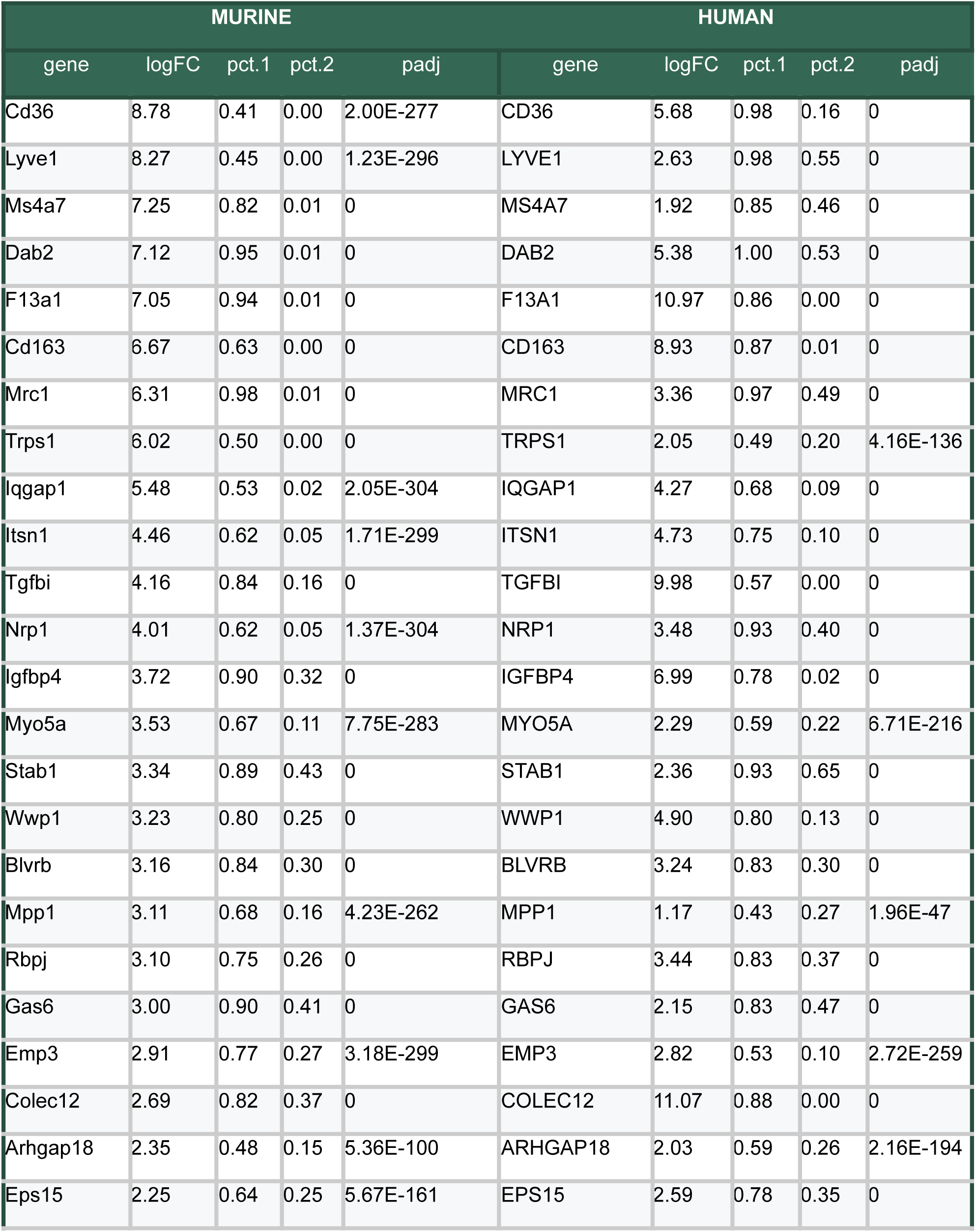
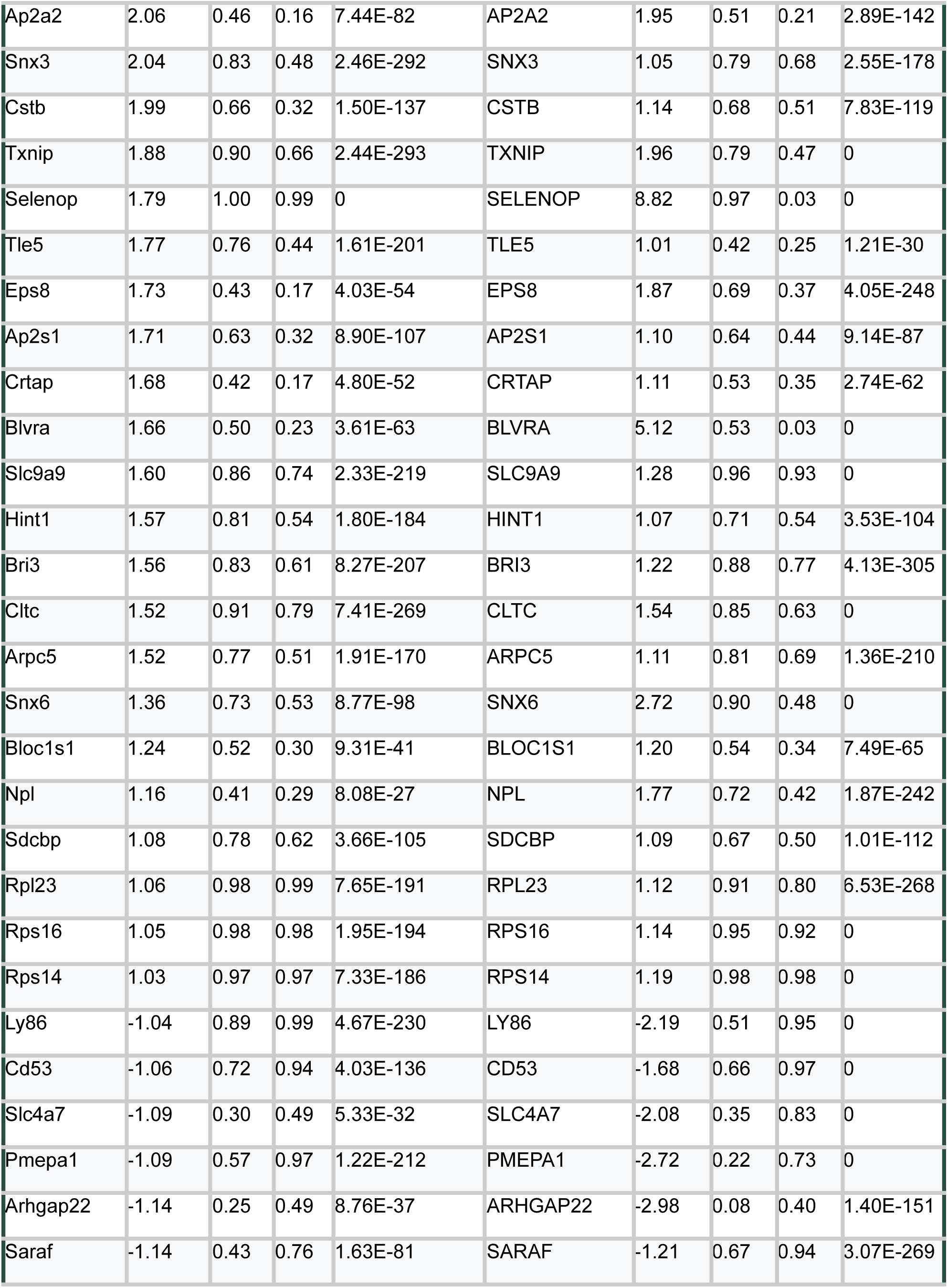
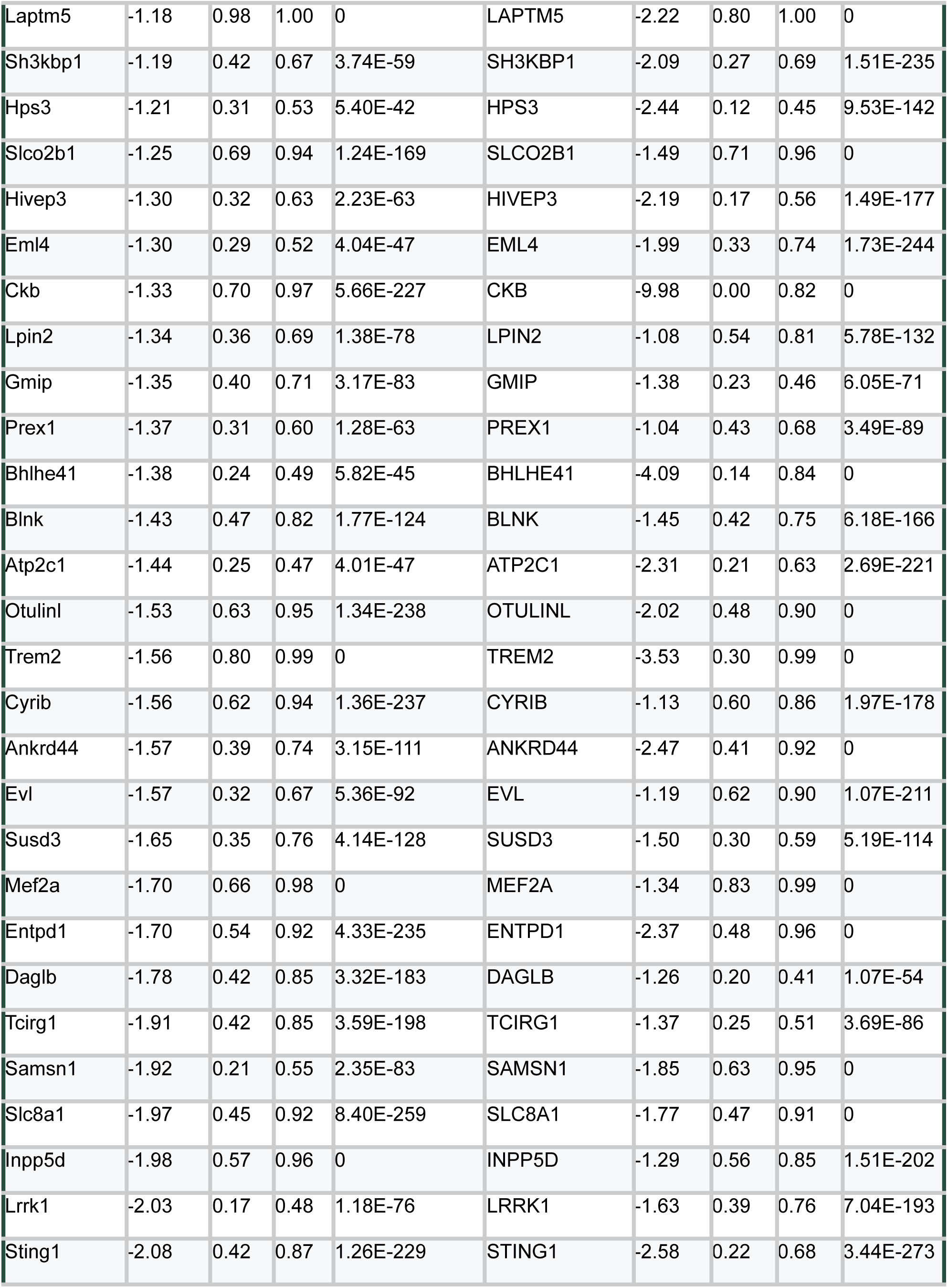
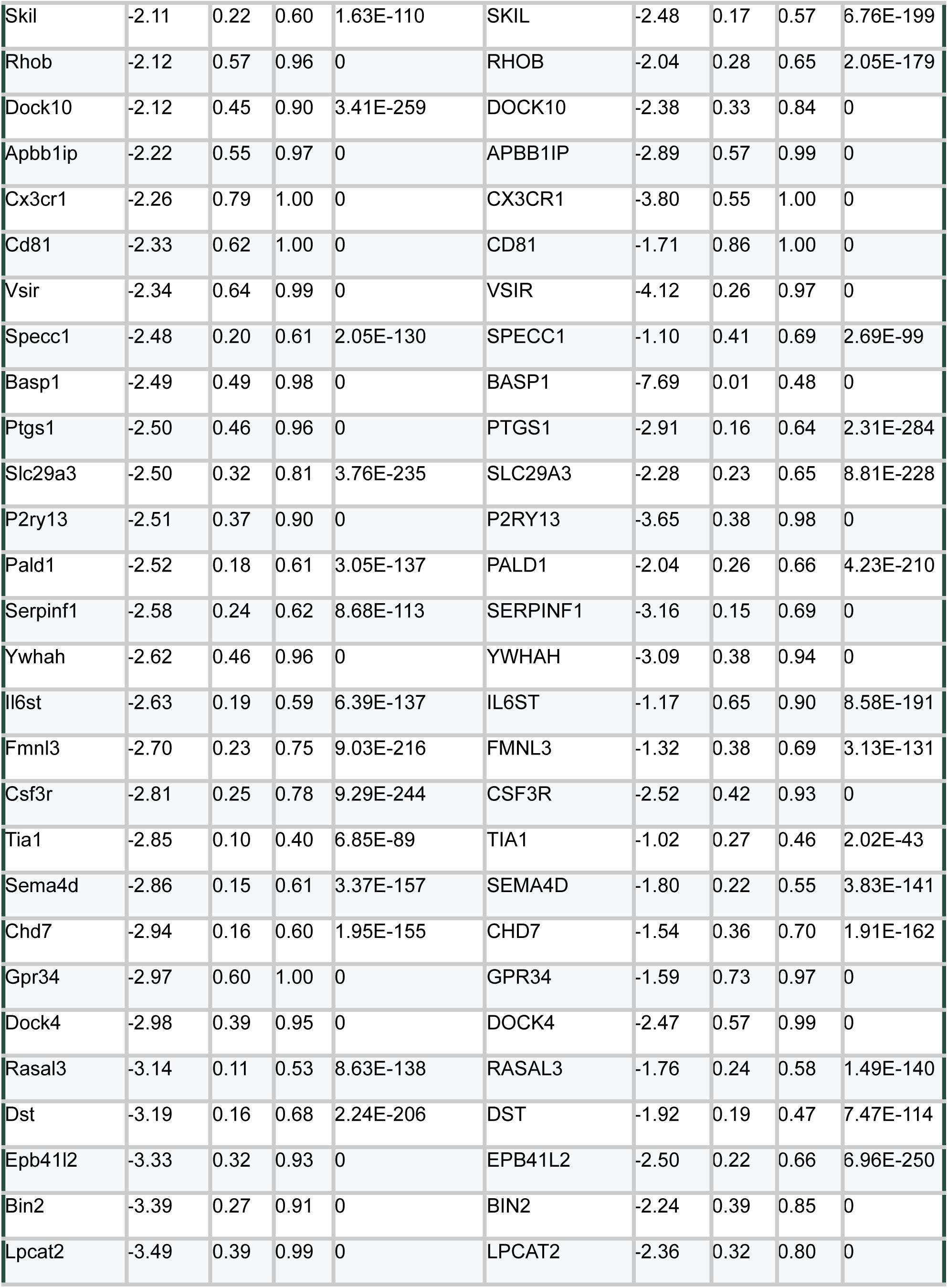
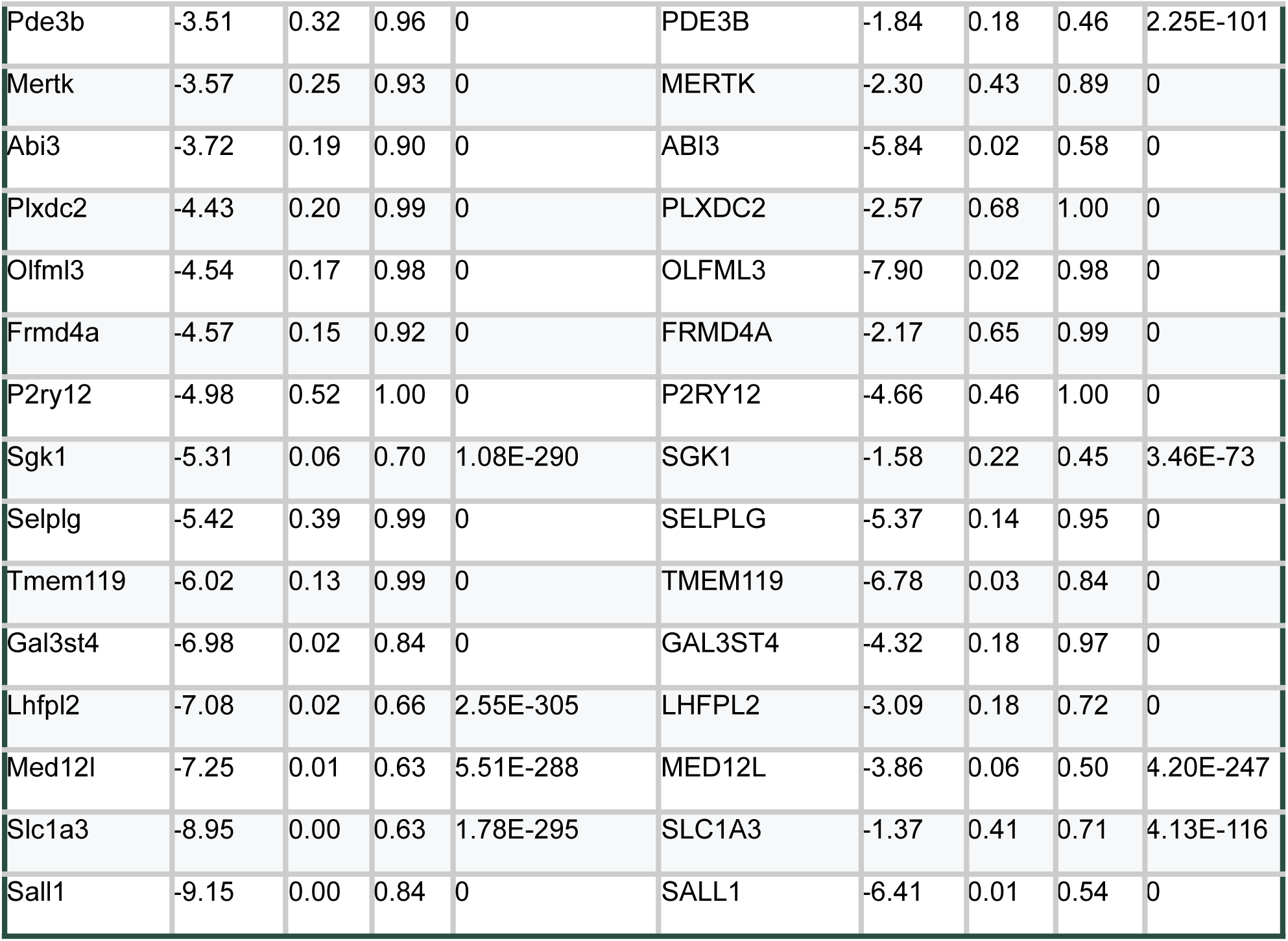
Conserved BAM versus microglia signature across human and murine origin. Gene set distinguishing BAMs from microglia of both human and murine origin. Genes were filtered based on a logFC threshold of 1 (corresponding to 2-fold enrichment or de-enrichment in BAMs), a min.pct of 0.4 (corresponding to at least 40% expression), and orthology verification across species.

**Table S3.** Flow cytometry antibodies for distinguishing BAMs and xBAMs. Recommended antibodies and dilutions for flow cytometry panel to identify BAMs and xBAMs in chimeric brains. For live cell preparations, we recommend identifying BAMs as muCD45^+^/ GR1^-^/ CD11b^+^CD64^+^/ CX3CR1^int^/CD38^+^ and xBAMs as huCD45^+^/huCD206^hi^. For fixed cell FEAST preparations, we recommend identifying BAMs as muCD45^+^/GR1^-^/CD68^+^/CX3CR1^int^/CD38^+^ (note: the panel is adjusted due to epitope loss following the fixed-cell protocol). The xBAM gating with huCD45^+^/huCD206^hi^ is compatible with the fixed cell FEAST protocol and does not require adjustment. See **Figure S3**.

| Antigen | Species reactivity | Fluorophore | Clone | Manufacturer | Cat # | Dilution | Note |
| --- | --- | --- | --- | --- | --- | --- | --- |
| CD45 | mouse | PE-Cy5.5 | 30-F11 | ThermoFisher | 35-0451-82 | 500 | Used 1:200 for FEAST |
| CD45 | mouse | BV605 | 30-F11 | BioLegend | 103155 | 500 |  |
| GR-1 | mouse | BUV805 | RB6-8C5 | BD Biosciences | 741920 | 100 |  |
| CD68 | mouse | APC-Cy7 | FA-11 | BioLegend | 137024 | 400 |  |
| CD206 | mouse | PE-Cy7 | C068C2 | BioLegend | 141720 | 200 |  |
| CD206 | mouse | BV421 | C068C2 | BioLegend | 141717 | 200 |  |
| CX3CR1 | mouse | BV711 | SA011F11 | BioLegend | 149031 | 800 |  |
| MHC-II | mouse | BV785 | M5/114.15.2 | BioLegend | 107645 | 200 |  |
| MBP | mouse/human/rat | AF647 | P82H9 | BioLegend | 850910 | 200 |  |
| CD38 | mouse | BUV395 | 90/CD38 | BD Biosciences | 740245 | 200 |  |
| CD64 | mouse | PE-Cy7 | X54-5/7.1 | BioLegend | 139313 | 100 |  |
| CD11b | mouse/human/rhesus | BV605 | M1/70 | BioLegend | 101237 | 200 |  |
| CD45 | human | AF488 | HI30 | BioLegend | 304017 | 200 |  |
| CD45 | human | BV421 | HI30 | BioLegend | 304032 | 200 |  |
| CD206 | human | PE | 15-2 | BioLegend | 321105 | 200 |  |
| CD206 | human | AF647 | 15-2 | BioLegend | 321116 | 200 |  |

